# Cerebellum Directly Modulates the Substantia Nigra Dopaminergic Activity

**DOI:** 10.1101/2022.05.20.492532

**Authors:** Samantha Washburn, Maritza Oñate, Junichi Yoshida, Jorge Vera, Ramakrishnan K. B., Leila Khatami, Farzan Nadim, Kamran Khodakhah

**Author notes:** Correspondence: Kamran Khodakhah.

## Abstract

Evidence of direct reciprocal connections between the cerebellum and basal ganglia has challenged the long-held notion that these structures function independently. While anatomical studies have suggested the presence of cerebellar projections to the substantia nigra pars compacta (SNc), the nature and function of these connections (Cb-SNc) is unknown. Here we show that the Cb-SNc form monosynaptic glutamatergic synapses with both dopaminergic and non-dopaminergic neurons in the SNc. Optogenetic activation Cb-SNc axons in the SNc rapidly increases SNc activity, elevates striatal dopamine levels, and increases the probability of locomotion. During ongoing behavior, Cb-SNc axons are bilaterally activated prior to ambulation and unilateral lever manipulation. The Cb-SNc axons show prominent activation to water reward, and higher activation for sweet water, suggesting that the pathway also encodes reward value. Thus, the cerebellum directly, rapidly, and effectively modulates basal ganglia dopamine levels and conveys information related to movement initiation, vigor, and possibly reward processing.

## Introduction

The effortless precision with which we can reach for and sip a glass of water or sign our name is perhaps most evident in patients who can no longer perform these tasks smoothly after a stroke, acute brain damage, or as a consequence of a neurodegenerative disorder. Understanding how the brain initiates and coordinates the precise motor sequences required for ordinary tasks, and how the process breaks down in the case of movement disorders, remains a vital goal of neuroscience. The brain generates, refines, and executes movements through a series of circuits involving the cerebral cortex, thalamus, cerebellum, and basal ganglia. The “loop” architecture of these circuits is well-described ^1–3^. The cerebral cortex sends extensive projections to the input areas of the cerebellum and basal ganglia, and the output nuclei of these regions are directed back via the thalamus to the same areas of cortex from which they receive input. The cerebellum and basal ganglia project to distinct, mostly non-overlapping regions of the ventrolateral and ventroanterior thalamus, respectively ^4^. For this reason, it had long been thought that subcortical interactions between the cortico-basal ganglia and the cerebello-thalamocortical circuits were limited and that, subsequently, the integration of basal ganglia and cerebellar outputs occurred in the cortex.

Recent anatomical evidence showing extensive subcortical connections between the cerebellum and basal ganglia in birds ^5^, rodents ^6^, and monkeys ^7–9^ suggests that there may be an alternative means of rapid coordination of basal ganglia and cerebellar outputs. Advances in viral tracing techniques have provided evidence for a disynaptic pathway from one of the output nuclei of the cerebellum, the dentate, to the striatum, the input nucleus of the basal ganglia, via the intralaminar nucleus of the thalamus ^6, 7^. It was later shown that this pathway enables the cerebellum to drive activity of neurons in the dorsolateral striatum with a very short latency, and to alter the sign of corticostriatal plasticity ^10^. A reciprocal pathway, from the subthalamic nucleus of the basal ganglia to the cerebellar cortex through the brainstem has also been anatomically described ^9^.

The striatum, however, is not the only target of direct subcortical projections from the cerebellum ^11^. Experiments performed almost 50 years ago provided functional ^12, 13^ and anatomical ^11, 14^ evidence of a pathway from the cerebellum to the substantia nigra pars compacta (SNc), a nucleus in the basal ganglia made up of primarily dopaminergic neurons that project to and modulate the striatum. The SNc is important for motor control, and neurons in this area are known to degenerate in Parkinson’s disease. It has been shown that strong trains of electrical stimulation, minutes long in duration, to the medial (fastigial), intermediate (interposed), and lateral (dentate) cerebellar nuclei can induce long-lasting changes in dopamine levels in both the SNc and the striatum ^12, 13^. Whether this is mediated directly, through a monosynaptic projection from the cerebellum to the SNc, or indirectly, via other brain regions, remains to be established.

Evidence in favor of a monosynaptic pathway from the cerebellar nuclei to the SNc comes from a number of separate studies using lesions, anatomical tracing, and imaging techniques. Degenerating terminals were observed in SNc after lesions to the fastigial cerebellar nuclei in the monkey ^15^, and to the interposed, dentate, and fastigial nuclei in the cat ^11^. The degenerating terminals appeared to be found specifically on putative dopamine neurons. More recently, cell-type specific, retrogradely-transported transsynaptic viral tracers have shown that the dentate nucleus of the cerebellum projects to SNc in the mouse ^14, 16^, while diffusion tensor imaging studies are consistent with the notion that a projection from the cerebellar nuclei to the SNc also exists in humans ^17, 18^.

While the studies described above have provided evidence that the cerebellum might be connected with the SNc, and that it can modulate activity of SNc neurons over the course of several hours, it is unclear whether the cerebellar nuclei can drive the activity of SNc neurons directly via a monosynaptic pathway. Given that SNc neurons are the main source of dopaminergic projections within the basal ganglia, if it is established that the cerebellum can rapidly modulate the activity of these neurons, the findings would have significant implications for how we understand motor control.

By combining optogenetics with electrophysiology, here we show that cerebellar nuclei neurons drive activity of SNc neurons via a monosynaptic, glutamatergic projection. The responses occur with latencies of a few milliseconds and are mediated by AMPA and NMDA receptors. Anatomical antero- and retrograde tracing experiments confirmed the presence of a monosynaptic projection from all three deep cerebellar nuclei to both dopaminergic and non-dopaminergic neurons in the SNc. Optogenetic activation of these projections, hereafter referred to as Cb-SNc projections, elevated dopamine levels in the striatum, and increased the probability that the mice walked. Using calcium as a proxy for neuronal activity, in fiber photometry experiments we found that the Cb-SNc axons bilaterally increased their activity just before the animal walked or performed a unilateral lever manipulation task. Intriguingly, in water-deprived mice, the Cb-SNc activity also increased when the mice consumed water, and the magnitude of the activity was greater for sweet water vs plain tap water. Thus, the cerebellum provides the SNc with information that might be relevant for initiation and possibly value or vigor of movement. Our findings prompt an earnest reevaluation of the circuits involved in motor coordination and provide a platform to test intriguing possibilities with regards to the nature of the information that is rapidly conveyed from the cerebellum to the dopaminergic nucleus of the basal ganglia.

## Results

### SNc neurons respond to stimulation of cerebellar fibers *in vivo*

We sought to determine whether neurons within the deep cerebellar nuclei (DCN) modulate the activity of the SNc neurons via a direct monosynaptic pathway. As a first approach to address this question, we used a combination of optogenetics and electrophysiology. In the first set of experiments, we examined whether optogenetic stimulation of cerebellar axons in SNc (Cb-SNc) could increase the activity of SNc neurons *in vivo.* Channelrhodopsin (ChR2) was expressed in DCN neurons, and an optrode was used to record from cells in the SNc, and to stimulate the ChR2 expressing cerebellar axons within the SNc (Figure 1A, Figure S1). The identity of the SNc neurons was determined based on the stereotaxic coordinates of the optrode at the time of the recordings, and post-hoc histological confirmation of recording sites (Figure S1D). We observed two types of responses to stimulation of Cb-SNc with short, 1 ms pulses of light: excitation, or excitation followed by a brief pause (Figure 1B, C). Approximately, two-thirds of the cells (n = 28/38 cells; N=17 mice) rapidly responded to stimulation of cerebellar inputs (Figure 1B). Of these cells, the majority (n= 26/28) responded with a transient increase in the number of spikes fired immediately following the stimulus (Figure 1B, C top panel), whereas in the remaining two the increase in the number of spikes was followed by a brief pause in firing (Figure 1B, C middle panel). Thus, optogenetic stimulation produced a large transient increase in the average firing rate (Figure 1D). To quantify the responses, we calculated the number of stimulation-evoked extra spikes by subtracting the number of expected spikes, given each cell’s baseline firing rate, from the number of spikes observed after the stimulus during the response (until the firing rate returned to within 3 standard deviation of the baseline). Overall, the average number of extra spikes was 0.64 ± 0.53 (Figure 1E), and the excitation phase of the response occurred with a short latency of 3.1 ± 2.9 ms (Figure 1F).

**Figure 1.**
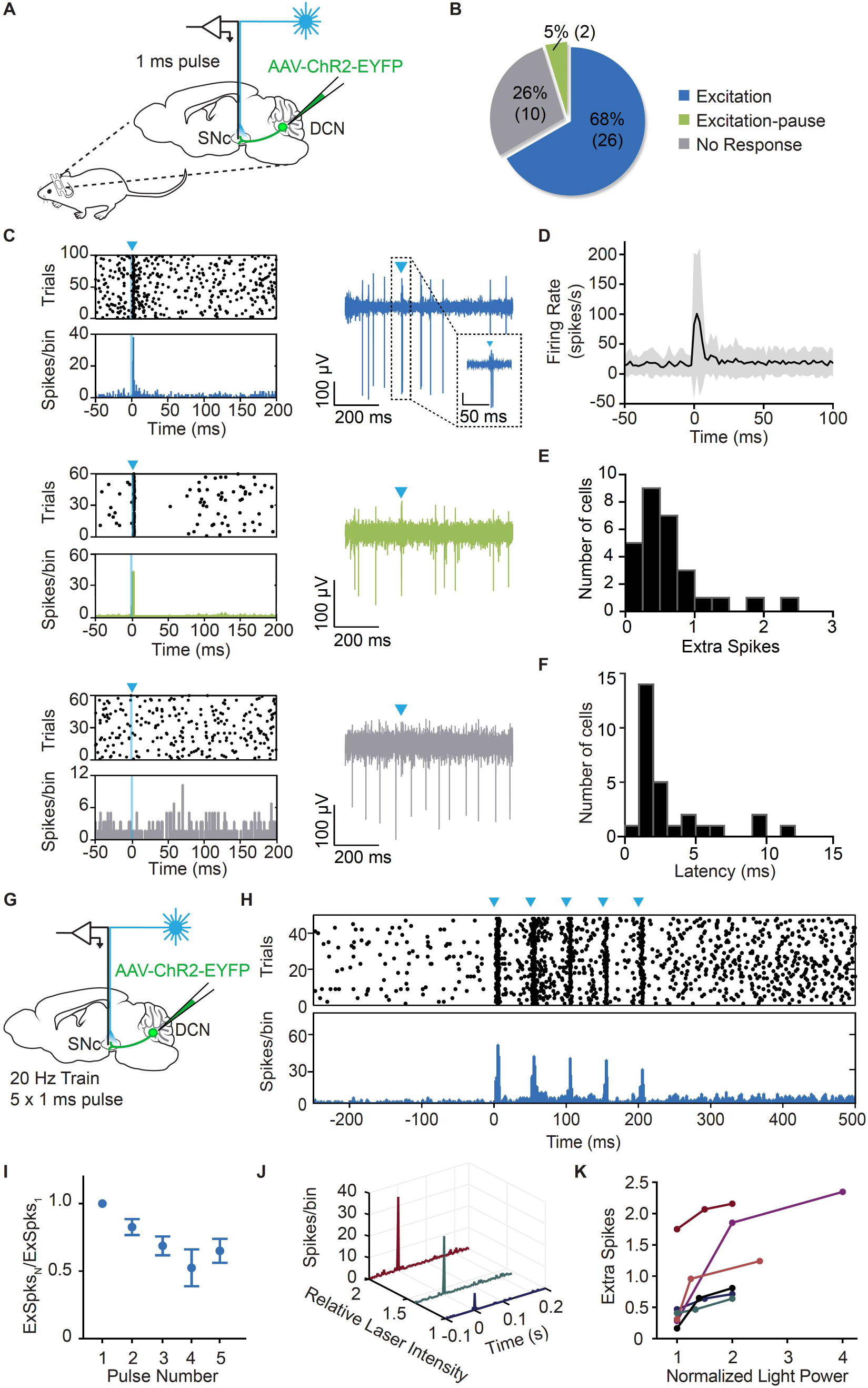
Neurons in SNc respond to activation of cerebellar fibers *in vivo*. (A) Mice were injected with Channelrhodopsin-EYFP (ChR2) in the DCN and an optrode was placed in the SNc to stimulate cerebellar fibers with 1ms pulses of light and record from SNc neurons while mice were head-fixed. (B) Proportion of responding cells in the SNc. From all cells recorded, approximately two thirds responded with excitation (blue) or excitation followed by a brief pause (green). The remaining third did not respond (grey). n = 38, N = 17. (C) Examples of a neuron that responded with excitation (top panel), excitation followed by a brief pause (middle panel), and no response (bottom panel). Left graph is raster plot (top) and PSTH (bottom), while right panel are example raw traces from each type of cell. Arrows indicate time of stimulation. (D) Average response of all responding cells. Shaded area is standard deviation. (E) Extra spikes histogram. Cerebellar stimulation evoked, on average, 0.64 ± 0.53 extra spikes. (F) Latency histogram. Cells responded to cerebellar stimulation with an average latency of 3.1 ± 2.9 ms. Mean ± SD. (G) Mice were injected with ChR2 in the DCN, and an optrode was placed in the SNc. Extracellular recordings were performed *in vivo* from neurons in SNc. Cerebellar fibers were stimulated with 5 pulses of light, 1 ms in duration at a frequency of 20 Hz. (H) Example raster plot (top) and PSTH (bottom) from a neuron that responded to a train of cerebellar stimulation. Arrows indicate time of stimulation. (I) Summary of cells recorded from SNc in response to train protocol (n = 6, N = 4). Response is extra spikes (ExSpks) evoked by each pulse normalized to the first pulse. Mean ± SEM. (J) The response of SNc neurons increased with increasing laser intensities. Example response of a neuron to increasing intensities of light stimulation. (K) Summary of cells recorded from SNc under stimulation with varying light intensities.

Cerebellar nuclei neurons are spontaneously active and are thought to convey information in their firing rates ^19^. We thus examined how neurons in the SNc responded to a train of stimuli, and to changes in the strength of the Cb-SNc inputs. We applied five, 1 ms pulses of light at a frequency of 20 Hz (Figure 1G) and found that SNc neurons increased their firing rate in response to each pulse in the train, with a modest adaptation in the amplitude of the responses (Figure 1H). The average decrease in the normalized response from the first to second pulse was 17.3 ± 14.4%, while the decrease in the normalized response from the first to last pulse was 35 ± 21.8% (Figure 1I). We then tested whether SNc neurons had larger responses when Cb-SNc axons were activated by light pulses of higher intensity and found this to be the case (Figure 1J). Doubling the light intensity increased the number of extra spikes by an average of 37.6 ± 15.5% (Figure 1K).

### Both dopaminergic and non-dopaminergic SNc neurons respond to stimulation of Cb-SNc projections *in vitro*

Next, we explored whether only dopaminergic neurons within the SNc receive input from DCN neurons because a previous study had suggested that the projection from the cerebellar nuclei to the SNc was specific to dopaminergic neurons ^11^. We injected animals with ChR2 in the DCN, and 3 weeks later acutely prepared horizontal brain slices for patch clamp recordings. We recorded from neurons in SNc in whole-cell, voltage-clamp mode (V_clamp_ = -60 mV), and stimulated Cb-SNc axons in the field of view through the objective using a LED (Figure 2A). The patch pipette contained 2 mM of Neurobiotin tracer to label each recorded cell, allowing for post-hoc immunohistochemical identification of TH-positive cells as a proxy for dopaminergic neurons (Figure 2B). The majority of the recorded cells (95%, n= 83/87 cells, N=23 mice) were TH-positive, while 5% of the recorded neurons (n= 4/87) were TH-negative. This ratio is consistent with the composition of dopaminergic and non-dopaminergic neurons that are thought to form the SNc ^20^, suggesting that we did not inherently bias these recordings to a specific cell type. Of interest, we found that both TH-positive (n= 41/83) and TH-negative (n= 2/4) cells responded to cerebellar stimulation (Figure 2C), suggesting that the projection is not cell-type specific.

**Figure 2.**
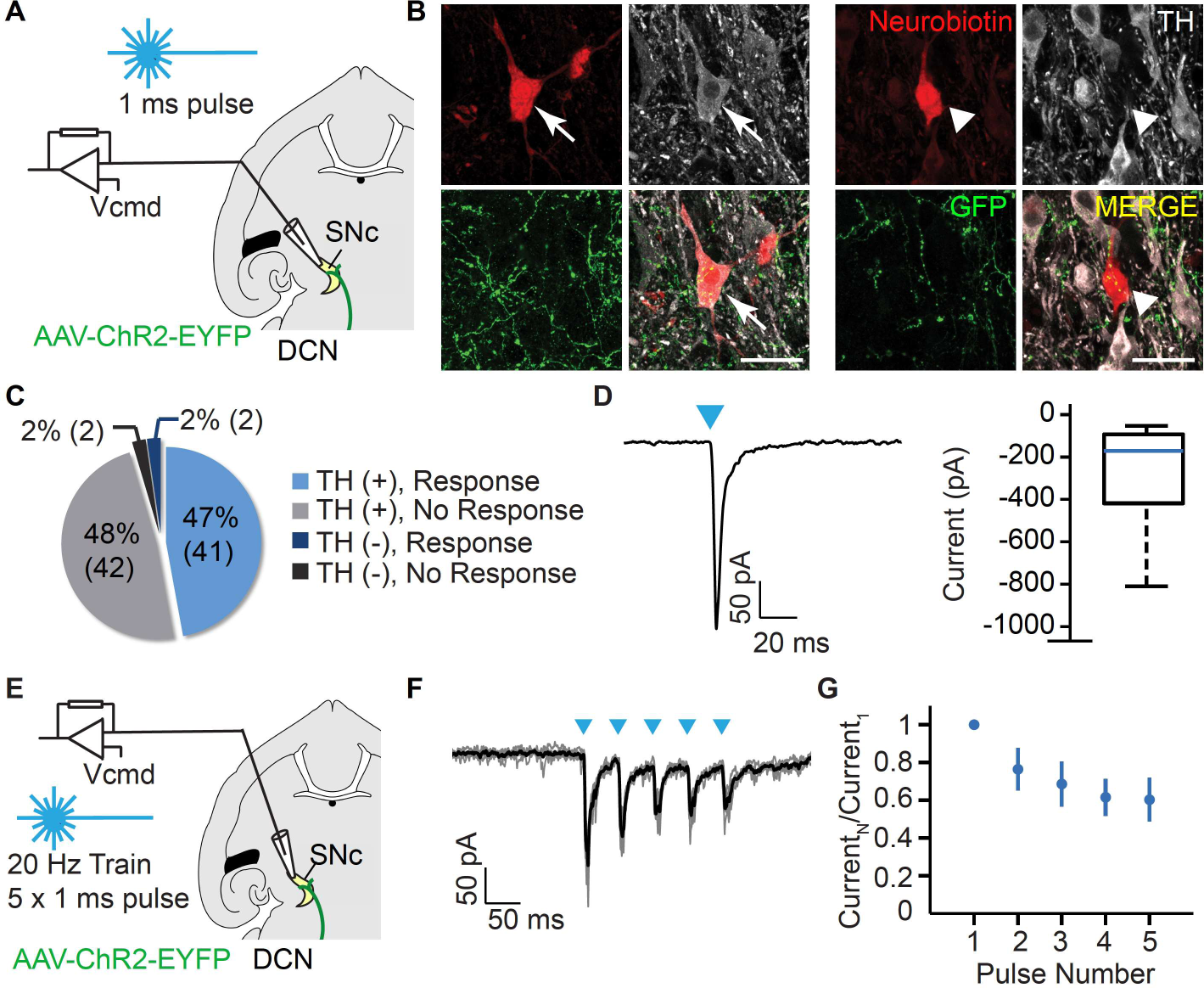
Dopaminergic and non-dopaminergic neurons in SNc respond to activation of cerebellar fibers in slices. (A) ChR2-expressing cerebellar axons were stimulated and neurons in SNc were recorded in whole-cell voltage clamp mode (Vhold = -60 mV) *in vitro*. (B) Recorded cells were filled with Neurobiotin for post-hoc histological analysis. Slices were stained for Neurobiotin (red) to identify recorded neurons and stained for TH (white) to characterize their phenotype. ChR2 cerebellar fibers (green) can also be seen with GFP in the field of view. (Left panel) Example of TH-positive cell recorded. (Right panel) Example of TH-negative cell recorded. Scale bars: 25 µm. (C) Proportion of responding cells in the SNc. The majority of recorded cells were TH-positive. Approximately half of all cells responded to cerebellar stimulation. These included both TH-positive and TH-negative cells. n = 87, N = 23. (D, left) Average of 5 sweeps from an example TH-positive cell that responded to 1 ms pulse of light stimulation. (Right) Boxplot showing distribution of current responses from TH-positive cells. (E) DCN neurons were infected with ChR2. Cerebellar fibers were activated with five 1-ms pulses of blue light at 20 Hz and SNc neurons recorded *in vitro*. (F) Response of a single neuron to train stimulus. Arrows indicate time of stimulation. (G) Summary of cells recorded from SNc in response to train protocol. Response is current evoked by each pulse normalized to the first pulse. n = 15, N = 10.

We recorded stimulus-evoked currents in SNc neurons in response to 1 ms light pulses, and found that, on average, the current evoked in the TH-positive neurons was -335.5 ± 417.4 pA (n = 41, N = 23) (Figure 2D). The two TH-negative cells responded with an average current of -76.3 and -123.2 pA, respectively.

As delineated earlier, SNc neurons respond to repeated *in vivo* Cb-SNc stimulation with a relatively modest adaptation. We thus examined the effect of repeated stimulation of Cb-SNc axons at a frequency of 20 Hz *in vitro* (Figure 2E). Consistent with our *in vivo* data, we found that trains of Cb-SNc stimulation reliably evoked responses in SNc neurons with each stimulation (Figure 2F). The amplitude of the response depressed by 31 ± 20% from the first to second pulse, and 45 ± 23% from the first to last pulse (Figure 2G).

### The projection from the cerebellum to the SNc is glutamatergic

The increase in firing rate observed in response to cerebellar stimulation *in vivo* suggested that the neurons projecting from the cerebellum to the SNc might be glutamatergic. Consistent with this possibility, when the cell was voltage clamped at +50 mV, the direction of the response was reversed, and the decay kinetics were prolonged suggesting the presence of N-methyl-D-aspartate (NMDA) receptors (Figure 3A, B). We examined this possibility using pharmacology. After identifying a responsive cell, NBQX was bath-applied to block α-amino-3-hydroxy-5-methyl-4-isoxazolepropionic acid (AMPA) receptors for glutamate, which mediate fast synaptic transmission. With V_clamp_ = -60 mV, we found that NBQX effectively blocked the evoked synaptic current (aCSF = 259.1 ± 109.2 pA vs. NBQX = 22.6 ± 15.7 pA, p = 0.0003, n = 8, N = 8) (Figure 3C). We then increased the holding potential to +50 mV to relieve the Mg^2+^ block of the NMDA receptors. We found that while the response at +50 mV persisted in the presence of NBQX, it was abolished by the NMDA receptor blocker D-APV (NBQX = 396.7 ± 238.7 pA vs. D-APV = 42.9 ± 10.3 pA, p = 0.0279, n = 5, N = 5) (Figure 3D), suggesting that both AMPA and NMDA receptors mediate the synaptic response.

**Figure 3.**
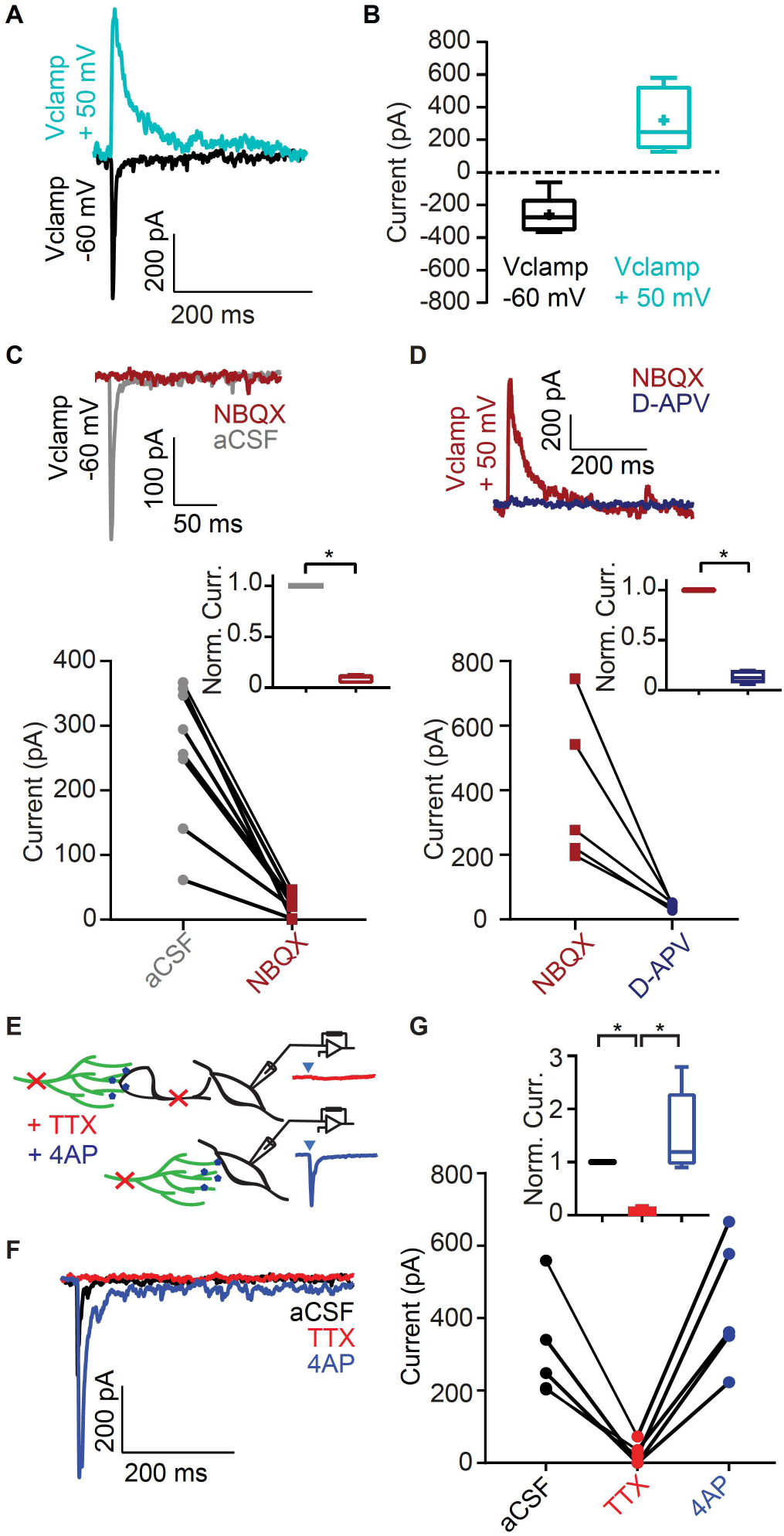
Cerebellar projection to SNc is glutamatergic and monosynaptic. (A) Example response to cerebellar stimulation at V_clamp_ = -60 mV and V_clamp_ = +50 mV. (B) Summary of responses at V_clamp_ = -60 mV (n = 8) and V_clamp_ = +50 mV (n = 5). (C) Example trace (top) and quantification (bottom) of responses of all cells in aCSF and NBQX at V_clamp_ = -60 mV. Absolute value of current is shown. Inset: Data normalized to aCSF, p = 0.0003, n = 8, N = 8. (D) Example trace (top) and quantification (bottom) of responses of all cells in NBQX and D-APV at V_clamp_ = +50 mV. Inset: Data normalized to aCSF, p = 0.0279, n = 5, N = 5. (E) Experimental schematic. TTX and 4AP were sequentially bath-applied. (F) Example traces of a cell in control condition (aCSF,), and after TTX (red) and 4AP (blue) treatment. (G) Quantification responses of all cells after TTX and 4AP. Absolute value of current is shown. Inset: Data normalized to aCSF; aCSF-TTX: p = 0.0174; TTX-4AP: p = 0.0098; aCSF-4AP: p = 0.2862; n = 5, N = 4.

### The projection from the cerebellum to the SNc is monosynaptic

The short latency of excitation observed in response to Cb-SNc stimulation *in vivo,* and the evoked synaptic currents *in vitro*, suggest that the projection from the cerebellum to the SNc might be monosynaptic. To directly test this possibility, we sequentially applied tetrodotoxin (TTX) and the voltage-gated potassium channel blocker 4-aminopyridine (4-AP) to our slices ^21^ (Figure 3E). Application of TTX blocks voltage gated sodium channels and prevents generation of action potentials. We found that bath application of 1 µM TTX abolished the response to stimulation of cerebellar axons (aCSF = 274.7 ± 161 pA vs. TTX = 24.19 ± 27.5 pA, p = 0.0189, n = 6). With TTX still in the bath, we applied 200 µM of 4-AP. 4-AP blocks low threshold voltage-gated potassium channel and allows for neurotransmitter release by direct optogenetic depolarization of nerve endings that express ChR2. We found that application of 4-AP restored the responses in all cases (4-AP = 374 ± 222.1 pA, p = 0.0198, n = 6), although as expected the kinetics of the responses were slower (Figure 3F, G). These data provide compelling evidence in support of the hypothesis that neurons in the SNc directly receive monosynaptic glutamatergic inputs from the DCN.

### Anatomical examination of cerebellar projections to the SNc

To complement our electrophysiological experiments delineated above, and to further characterize the nature of the connectivity between the cerebellum and the SNc, we performed a combination of viral-mediated anterograde and retrograde tracing experiments.

We first took advantage of an EGFP reporter mouse, RCE:loxP, in which expression of Cre drives expression of the fluorescent protein EGFP. It is known that the adeno-associated virus AAV1 has anterograde transsynaptic spread properties and can drive expression of Cre in postsynaptic neuronal targets. As such, when injected into a specific brain region, it first infects the cells in that region, and then spreads to infect cells that are monosynaptically targeted by the neurons in that region ^22, 23^. We injected AAV1-hSyn-Cre bilaterally into the DCN (Figure 4A). As expected, cells in the DCN that were infected by the virus expressed Cre-dependent EGFP (Figure 4B), as well as cells that received monosynaptic inputs from the DCN. Examination of the SNc showed many neurons that were fluorescent and therefore monosynaptic targets of the DCN (Figure 4C). We used TH staining to determine whether the neurons that received direct DCN inputs were dopaminergic or non-dopaminergic. We found that both TH-positive and TH-negative cells received cerebellar inputs, consistent with the observations reported earlier based on the *in vitro* electrophysiological experiments (Figure 4C, right panel).

**Figure 4.**
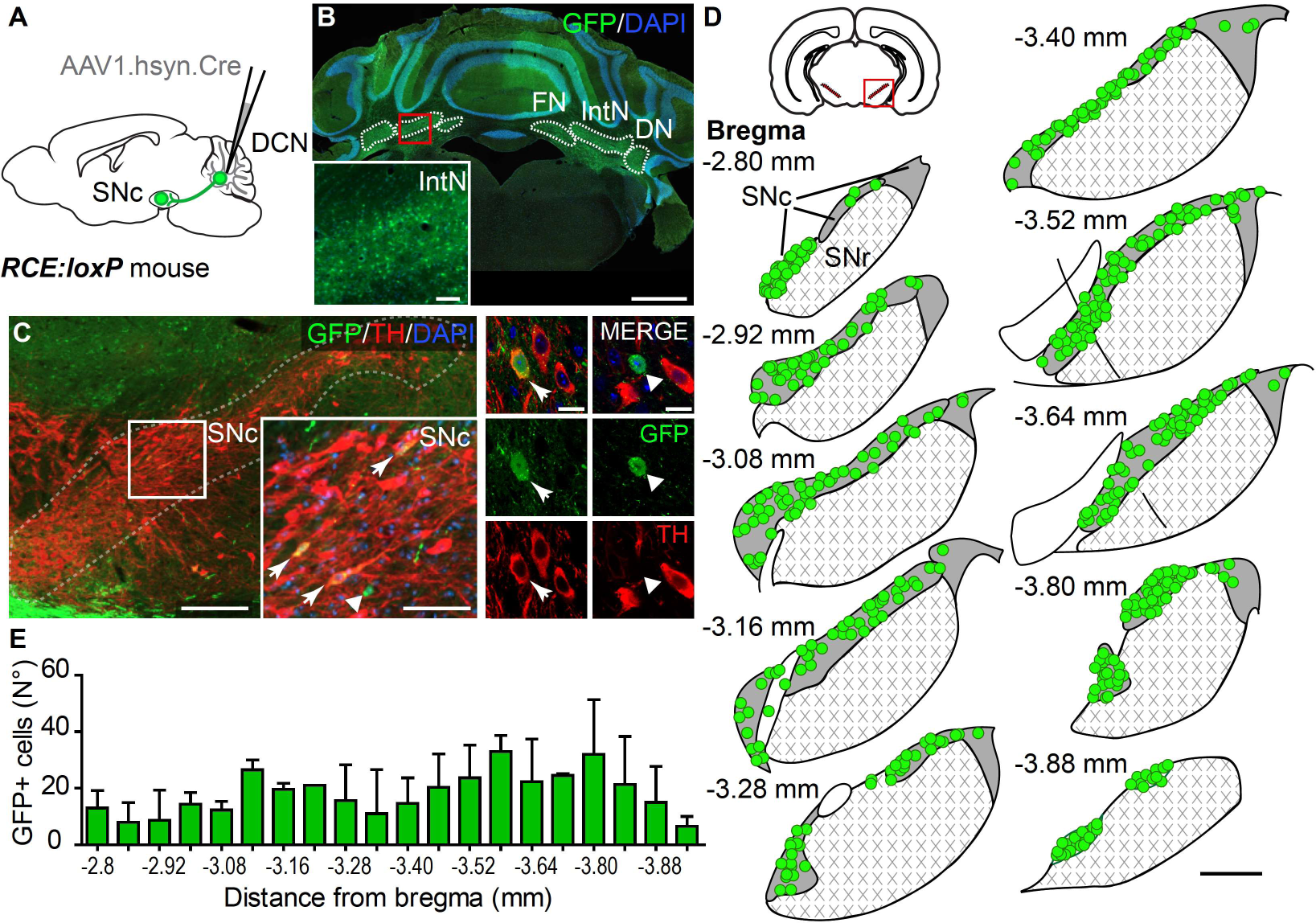
Anatomical evidence of monosynaptic cerebellar projection to the SNc. (A) Experimental design. AAV1-Cre was injected in the DCN of RCE:loxP mice to trace DCN-recipient cells in the SNc. (B) Representative image of cerebellum showing bilateral injection in the DCN. AAV1-Cre virus is visualized with GFP in green. DAPI staining is shown in blue. DN: dentate nucleus, IntN: interposed nucleus, FN: fastigial nucleus. Scale bar: 750 µm. Zoom in of the infected cells in the interposed nuclei. Scale bar: 100 µm. (C) (Left panel) Representative image of the SNc showing DCN-recipient cells (green). TH staining was performed to delimit the structure and to label for dopaminergic neurons (red). Scale bar: 100 µm. Zoom in of the SNc showing DCN-recipient cells. Arrows point TH-positive and arrowhead, TH-negative GFP cells. Scale bar: 50 µm. (Right panel) Examples of dopaminergic (left column) and non-dopaminergic (right column) DCN-recipient cells. Scale bars: 25 µm. (D) Distribution of individual DCN-recipient neurons along the antero-posterior (A-P) SNc. (Top left) Schematic of brain region analyzed. All DCN-recipient cells were mapped in representative bregma schematics from anterior to posterior SNc. Only right SNc is shown. Cells of N = 3 mice were mapped. Scale bar: 250 µm. SNr: substantia nigra reticulata. (E) Quantification of the number DCN-recipient cells in the SNc along the A-P axis (represented by bregma position). Mean ± SD, N= 3.

To delineate the SNc regions that receive direct projections from the DCN, we analyzed serial sections encompassing the entire SNc to map the distribution of DCN-targeted cells (Figure 4D). Quantification of target cells in antero-posterior coronal serial sections revealed that the DCN targeted the entire SNc with no major regional variations in distribution (Figure 4D, E). Thus, the DCN sends monosynaptic projections to dopaminergic and non-dopaminergic neurons throughout the SNc.

We then used retrograde tracing of inputs to the SNc to delineate whether neurons from only one particular cerebellar nuclei send projections to the SNc, or alternatively, whether the projections arise from all three nuclei. To do so, we injected a retroAAV virus expressing Cre in the SNc of RCE mice^24^. To label the site of injection we co-injected the retroAAV-Cre with an AAV expressing tdTomato (Figure 5A, B). Analysis of cerebellar sections showed that DCN projections arise from neurons in all three nuclei; the Dentate (DN), Interposed (IntN) and Fastigial (FN) nucleus (Figure 5C). However, a larger fraction of the projections originated from the IntN and the DN compared to FN (Figure 5D). Mapping and analysis of distribution of DCN projection neurons along the antero-posterior axis showed that the SNc receives cerebellar input from most of DN and IntN (Figure 5E). However, in the FN, the neurons were enriched in the ventro-posterior region (Figure 5E, right column), similar to what has been reported previously ^14, 25^.

**Figure 5.**
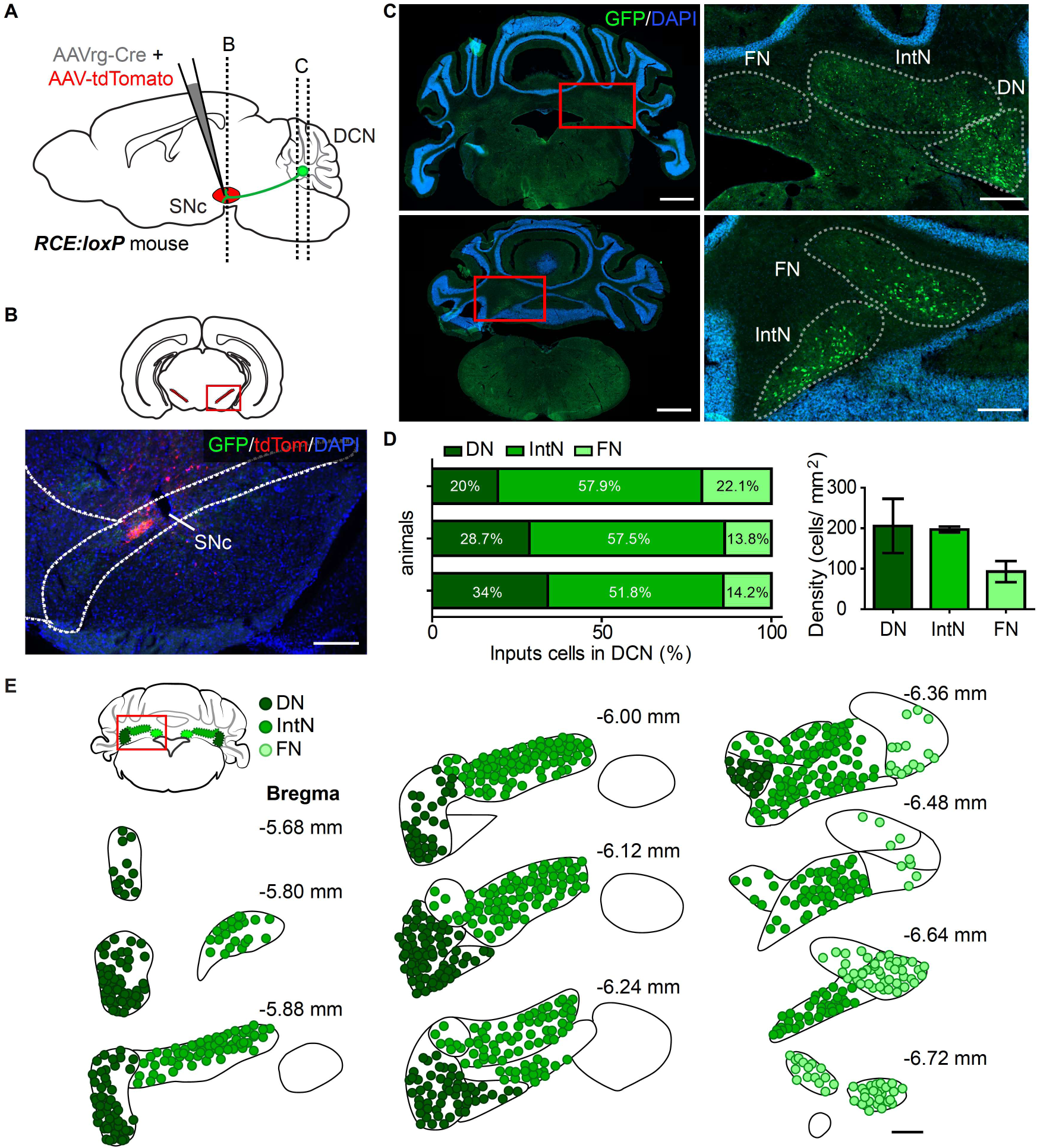
All three nuclei of the DCN project to SNc. (A) Experimental design. retroAAV-Cre was injected in the SNc of RCE:loxP mice to trace DCN input cells. An AAV expressing tdTomato was co-injected in the SNc to label the site of injection. B and C dash lines show bregma position images in B and C. (B) Scheme showing the region of analysis in the SNc. Representative image of the SNc showing site of injection in red (tdTomato). DAPI was stained in blue. Scale bar: 100 µm. (C) Anterior and posterior examples (top and bottom left, respectively) of cerebellar sections showing input cells in the DCN. Scale bar: 1 mm. (Right) Zoom in showing input cells in different cerebellar nuclei in green (GFP). Scale bar: 250 µm. (D) (Left) Percentage of input cells in each nucleus of the DCN per mouse. (Right) Density of input cells per nuclei. Mean ± SD, N= 3. (E) Distribution of input cells along the antero-posterior DCN. (Top left) Schematic of brain region analyzed. All DCN-input cells were mapped in representative bregma schematics from anterior to posterior DCN. Cells in each nucleus are colored in different green tones. Only left DCN is shown. All cells of one mouse are shown. Scale bar: 250 µm.

### Activation of cerebellar axons in the SNc drives dopamine release in the striatum

The data presented so far demonstrate that monosynaptic cerebellar projections can drive activity in dopaminergic and non-dopaminergic neurons in the SNc. Since activity of SNc dopaminergic neurons regulates dopamine levels within the striatum, we explored whether optogenetic activation of Cb-SNc axons could increase dopamine levels in the striatum. To measure dopamine, we expressed the fluorescent dopamine sensor dLight1.1 in striatal neurons ^26^, and bilaterally injected ChR2 in the cerebellar nuclei. A fiber optic was implanted to target SNc to allow for the optogenetic activation of Cb-SNc axons, and a second fiber optic was positioned in the ipsilateral striatum to measure changes in dopamine levels. The mouse was head restrained on a wheel that allowed it to freely ambulate, and the speed of the wheel movement was measured (Figure 6A, Figure S2).

**Figure 6.**
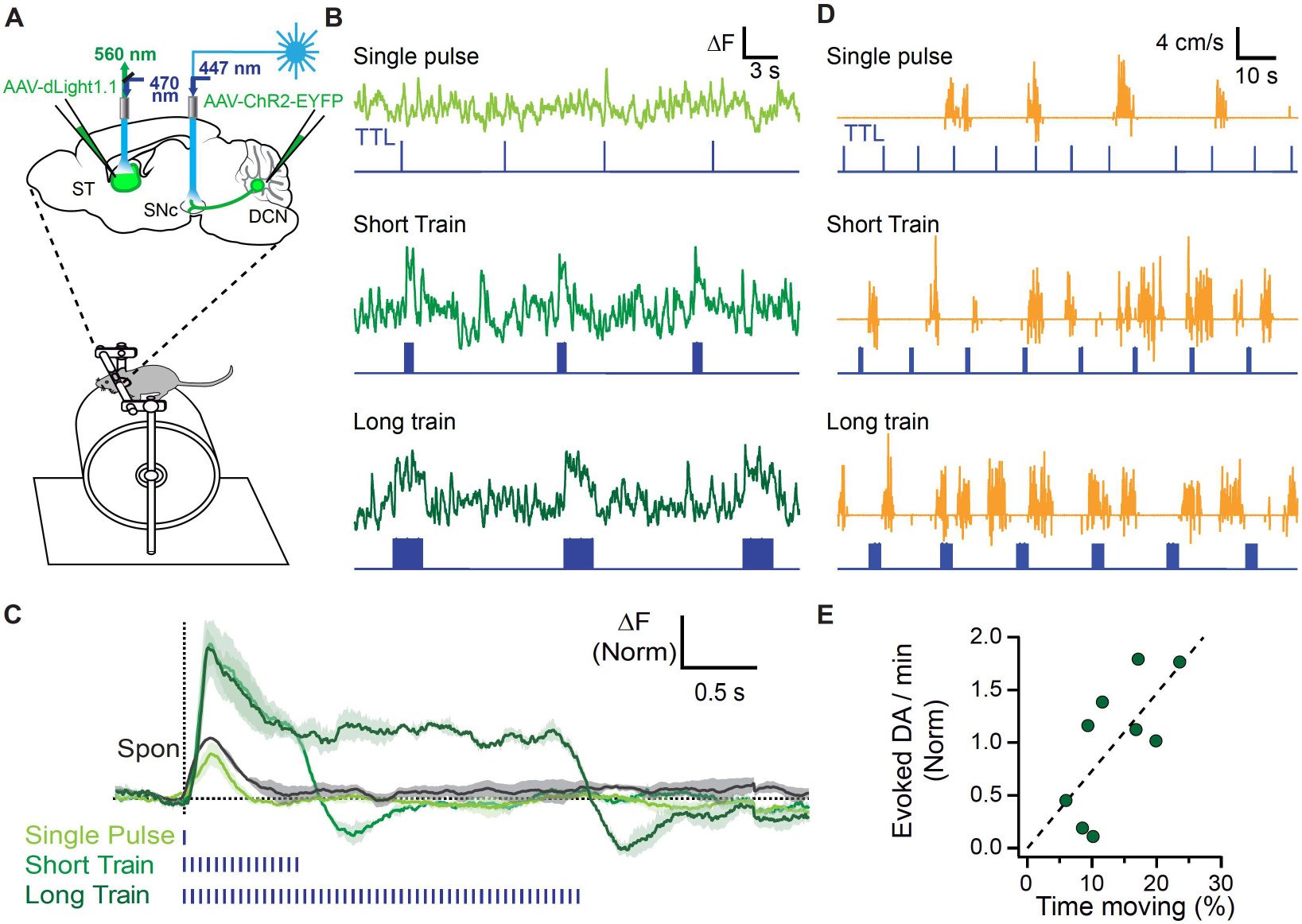
Optogenetic activation of Cb axons in SNc evokes dopamine release in the striatum of behaving mice. (A) Mice expressing channelrhodopsin in the DCN and the dopamine sensor dLight1.1 in the striatum where head-fixed to a metal frame on top of a movable cylinder that allows free walking. Cb-SNc axons were excited through a fiber optic implanted in SNc while dopamine fluctuations were recorded with a fiber optic implanted in the striatum. (B) Fiber photometry signals recorded from the striatum (green) during optogenetic stimulation of Cb-SNc axons (470 nm, 1 mW; TTL signal in blue) with single pulses, and short or long trains (1 ms pulses at 20 Hz). (C) Stimulus-triggered average of striatal dLight1.1 signals (mean ± SE, N=3) evoked with optogenetic activation of Cb-SNc axons. Spontaneous fluctuations (Spon) were independent of stimulation, and aligned to the onset of each event. (D) Wheel speed at different patterns of Cb-SNc axons activation with optogenetics. Labels as in B. (E) Correlation of evoked dopamine and locomotor activity as a function of Cb-SNc axons stimulation. Pearson’s linear correlation, r=0.7, P = 0.035 (N=3). Data shown are evoked dopamine per minute (integral of fluorescence after stimulus onset normalized by the mean value for each animal) vs. the percentage of time spent walking during the time of Cb-SNc axons stimulation with single, short or long train.

Under baseline conditions, we found that striatal dopamine levels showed fast transient spontaneous fluctuations of about 200 ms in duration (Figure 6B, C). In the same animals, brief 1 ms optogenetic activation of Cb-SNc axons reliably produced transient increases in the striatal dopamine levels (Figure 6B, Single pulse). These cerebellar-evoked dopamine transients were comparable in amplitude and kinetics to the spontaneous dopamine fluctuations seen in the periods without stimulation (Figure 6C). Given that DCN neurons are continuously active, we explored the impact of repeated stimulation of the cerebellar axons with short (15, 1 ms pulses at 20 Hz), and long (50, 1 ms pulses at 20 Hz) trains on striatal dopamine levels (Figure 6B, C). These trains demonstrated that repeated activity of Cb-SNc inputs can generate a sustained increase in the striatal dopamine levels.

It has been proposed that striatal dopamine levels contribute to movement initiation and vigor ^27–32^. We thus explored whether activation of the nigrostriatal circuit by cerebellar inputs could promote movement. To do so, we optogenetically activated the Cb-SNc axons to varying degrees and measured the time that mice walked on the wheel as a fraction of the total time. We found a positive correlation between the dopamine signals evoked with Cb-SNc axon stimulation, and the fraction of locomotion time, demonstrating that activation of this pathway could increase the probability of movement (Figure 6D, E; r= 0.7, P = 0.035). While these data suggest that the cerebellar projections to the SNc can promote movement, for the pathway to have a meaningful contribution under physiological conditions, it would be expected that these cerebellar axons would be active as the animal moves. To examine the validity of this prediction, we expressed a green calcium sensor, GCaMP7, in cerebellar axons by injecting the virus into the DCN, and a red calcium sensor, jRGECO1, in the SNc. This configuration allowed us to use changes in the intracellular calcium as a proxy for neuronal activity, and to simultaneously monitor the activity of the SNc neurons, and Cb-SNc axons, using a single fiber optic as the mouse walked on a wheel at will (Figure 7A, Figure S3).

**Figure 7.**
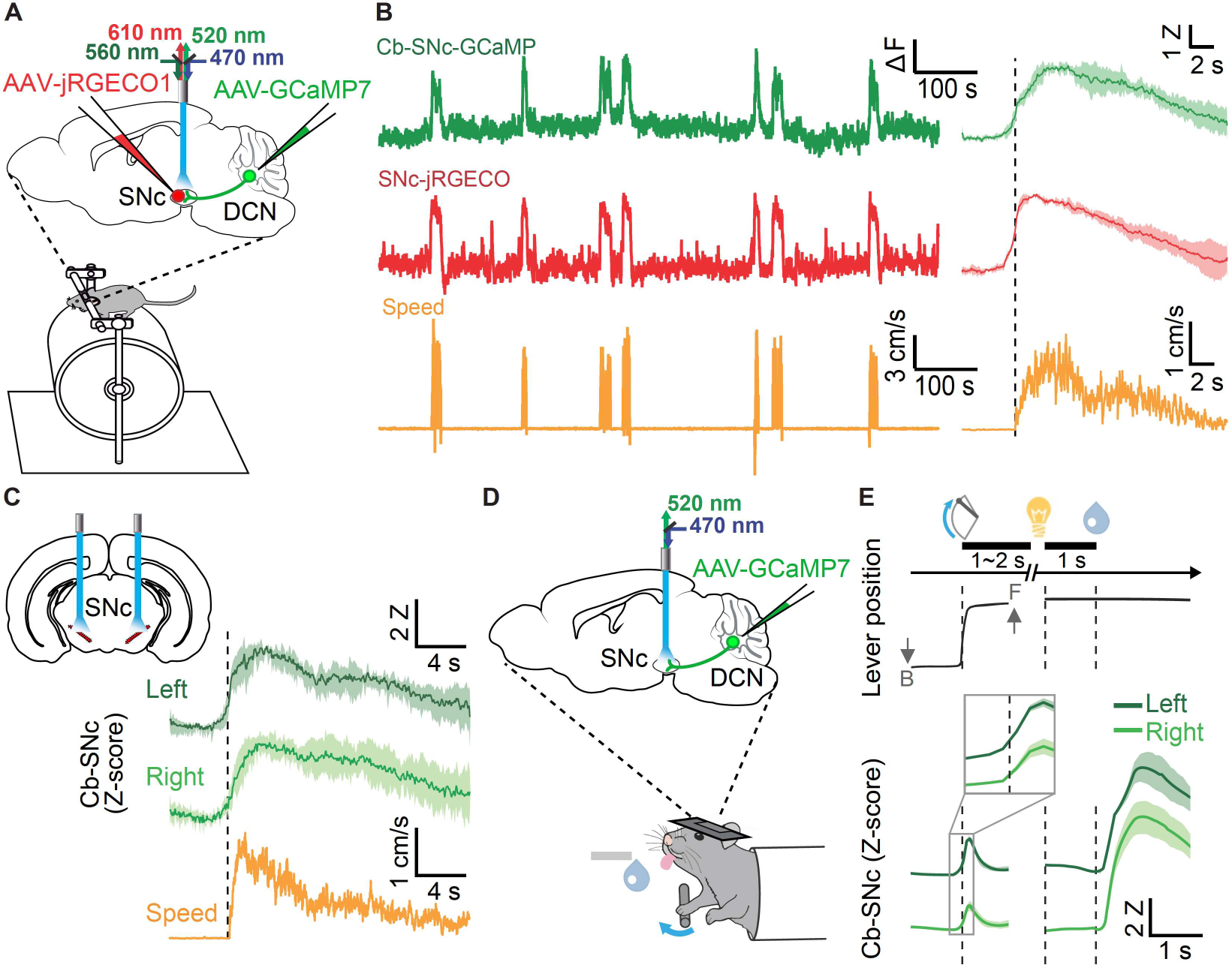
Natural activation of Cb axons in SNc during simple behaviors. (A) Mice expressing complementary calcium indicators in the Cb (GCaMP7) and the SNc (jRGECO1) were head-fixed to a metal frame on top of a movable cylinder that allows free walking. Fluorescence from Cb-SNc axons and SNc neurons were recorded simultaneously (at alternated frames) through the same fiber implanted in SNc. 1.6 mW/mm^2^ 470nm- and 3.2 mW/mm^2^ 560nm-LED were used for GCaMP and jRGECO signal measurement. (B) (Left) Fiber photometry signals Cb-SNc axons (Cb-SNc-GCaMP) and SNc neurons (SNc-jRGECO), together with the speed of the wheel (Speed), during spontaneous locomotor activity. Traces at the right show the average activity of each calcium sensor associated to locomotor events onset (dotted line) (mean ± SEM, N=4). (C) Bilateral photometry recordings of Cb-SNc axons expressing GCaMP showing activity associated to onset of locomotor events (dotted line) (mean ± SEM, N=4). (D) Schematic of photometry recordings of Cb-SNc axons during lever-manipulation task. AAV-expressing GCaMP7 was injected in the DCN and optic fiber implanted in the SNc. 1.6 mW/mm^2^ 470nm-LED was utilized for GCaMP signal measurement. Schematic of the task is shown at the bottom. Head-restricted mice were trained to push a lever to receive water as reward, between 1-2 seconds after pushing the lever, a light cue appears, and then 1 second later the water is delivered. (E) Mean of the lever position (Top), and Z-score of Cb-SNc axons activity (bottom) across the task. Calcium signals were measured bilateral in left (dark green) and right (light green) SNc. The data is aligned at lever push initiation (first dashed line), light cue (second dashed line) and reward delivery (third dashed line). B and F represent backward and forward position of the lever, respectively. The inset indicates Cb-SNc activity aligned at lever push initiation with ± 200 ms time window. Lever position, N = 4 mice x 5 sessions. Right Cb-SNc activity, N = 5 sessions x 4 sites (from 4 mice). Left Cb-SNc activity, N = 5 sessions x 3 sites (from 3 mice). Time-bin duration is 10 ms in lever position and 60 ms in neuronal activity. Z means Z-score (see Methods). Mean ± SEM.

When mice were not walking on the wheel, calcium signals from both the Cb-SNc axons and SNc showed a stable baseline, with occasional low amplitude transient increases (Figure 7B, left panel). However, every time that the mice engaged in locomotor activity, there was a prominent increase in the activity of the SNc, and the cerebellar axons within it (Figure 7B, left panel). The duration of both signals correlated with the length of the time that the mouse walked. Moreover, the alignment of calcium signals relative to the onset of wheel movement demonstrated a tight temporal correlation between the increase in the activity of SNc and Cb-SNc axons and the onset of locomotor activity (Figure 7B, right panel). Simultaneous bilateral recordings demonstrated that, with ambulation, the increase in the activity of the cerebellar axons in the SNc was similar in its time course and amplitude in the two hemispheres (Figure 7C).

The increase in the activity of SNc neurons, and the Cb-SNc axons appeared to start prior to the onset of the wheel movement (Figure 7B, C). However, walking is a complex motor task that engages a wide array of muscles, and the animals often showed other movements, such as whisking or tail flicks that preceded ambulation. It was not practical to accurately quantify these movements with our experimental setup. Thus, we cannot rule out that these movements could be the cause of, or a major contributing factor to, the increase in the activity of SNc and Cb-SNc prior to onset of wheel movement. Similarly, the complexity of the movements associated with walking, and the fact that muscles on both sides of the body are engaged during the process, might account for the fact that the signals on both hemispheres were comparable.

To explore the temporal correlation between the changes in the activity of Cb-SNc and the onset of movement in more detail, and to establish whether the cerebellar signals are unilateral or bilateral with unilateral movement, we trained a second cohort of mice to perform a simple lever-manipulation task. In this task, mice were trained to move a lever forward using their right paw only. To aid with the training, the mice were partially water deprived, and were given water as reward when they completed the task. Thus, the paradigm consisted of the mouse initiating the task by pushing a lever forward. Then, after a random delay of 1-2 seconds a light cue was delivered, and 1 second later, a small amount of water was dispensed as reward. Mice were injected in the DCN with an AAV-expressing the calcium sensor GCaMP7, and fiber optics were implanted bilaterally in each hemisphere of the SNc to measure Cb-SNc activity while mice perform the task (Figure 7D, Figure S4). Bilateral recordings of the calcium signals from the Cb-SNc axons showed that, concurrent with the unilateral movement of the right paw as the mouse pushed the lever forward, cerebellar axons in the SNc of both hemispheres increased their activity (Figure 7E), similar to the bilateral signals seen with ambulation. Moreover, the neuronal activity in the Cb-SNc axons started to increase prior to the onset of lever movement (Figure 7E, inset). As mentioned, to train the animals to perform the lever manipulation task they were partially water deprived and received a small amount of water as reward. Intriguingly, much larger signals were detected bilaterally in the Cb-SNc axons when the animals received the water reward compared to when they pushed the lever (Figure 7E).

### The activity of cerebellar axons in the SNc contains reward-related information

The large bilateral signals in the Cb-SNc when the mice received reward were unexpected and demanded closer scrutiny. We thus switched to a simple Pavlovian task where partially water-deprived mice were given water as reward one second after a light cue. As in the previous experiment, mice were injected with GCaMP7 in the DCN a bilaterally implanted with fiber optics in both hemispheres of the SNc to monitor the activity of Cb-SNc axons (Figure 8A). To explore the nature of the Cb-SNc signals, in random trials the tap water (Regular reward) was either omitted or replaced with either the same volume of 0.1% Saccharine water (High reward), or triple dispensation of regular water (3 x Regular at 250 ms intervals; Figure 8B). There was a prominent increase in the Cb-SNc signals every time that the mice received reward, but not when reward was omitted (Figure 8B). The amplitude and duration of the Cb-SNc signal was greater when the mice received the High reward sweet water than when they received the Regular tap water reward. When the tap water was delivered three times in succession, the Cb-SNc signal was longer, and individual peaks that correlated with the dispensation of each drop of water were clearly discernable (Figure 8B). The Cb-SNc signals in this task, too, were bilateral (Figure 8C).

**Figure 8.**
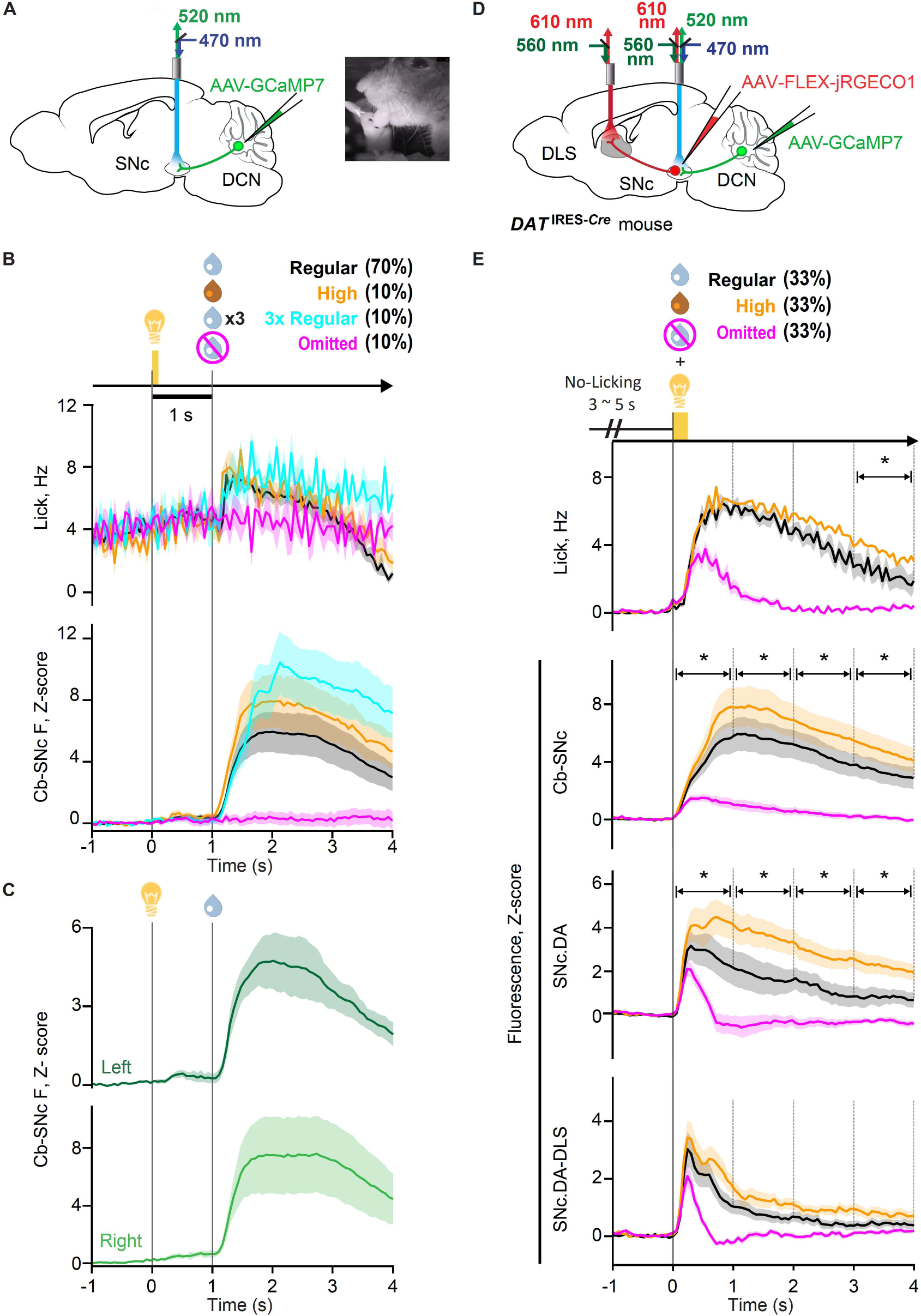
Reward evaluation in the Cb-SNc activity. (A) (Left) Design of the single-color fiber photometry experiment. Calcium sensor (GCaMP7) was injected in the DCN. Optic fibers were implanted in the SNc to record Cb-SNc axons. 1.6 mW/mm^2^ 470nm-LED was utilized for GCaMP signal measurement. (Right) Image of a mouse licking liquid rewards from a spout. This spout has two ports delivering different types of liquid reward separately. (B) Average of licking frequency and Z-scored of Cb-SNc activity during the task. Mice were given a reward (drops) 1 second after a light cue (yellow light bulb). Reward was Regular, 4 µL tap water (black). High, 4 µL 0.1% saccharine water (yellow). 3xRegular, three shots of the 4 µL tap water (cyan). Omitted, no reward (magenta) with a high or low probability. Data is aligned to the cue and reward delivery. Licking frequency, N = 4 mice. Cb-SNc activity, N = 7 sites. Mean ± SEM. (C) Average of Cb-SNc activity in the right (light green) and left hemisphere (dark green) across animals. Z-scored activity is aligned to the cue and reward delivery. Right Cb-SNc activity, N = 4 sites (from 4 mice). Left Cb-SNc activity, N = 3 sites (from 3 mice). Mean ± SEM. (D) Design of the dual-color fiber photometry experiment. AAV-expressing GCaMP7 was injected in the DCN while Cre-dependent jRGECO was injected in the SNc of DAT-Cre miceo, and optic fibers were implanted in the SNc and dorsolateral striatum, DLS. 1.6 mW/mm^2^ 470nm- and 3.2 mW/mm^2^ 560nm-LED were utilized for GCaMP and jRGECO signal measurement, respectively. (E) Mice were trained to stop licking between 3-5 seconds prior the cue and reward delivery. Probability of reward or omitted trial is described. Average of licking frequency and neuronal activity (Z-score) at the cue onset across animals and recording sites in Cb-SNc axons, SNC.DA somas at the SNc and SNC.DA-DLS axons. Asterisks indicate significant difference between Regular and High reward trials in 1 sec time-bins (p < 0.05 for post-hoc Holm-Sidak test following one-way RM-ANOVA). Licking frequency: N = 8 mice; neuronal activity in each region: N = 16 sites (bilateral recordings from 8 mice); signals recorded at 60 ms sampling intervals; data shown as mean ± SEM.

Consumption of water requires licking, a motor task, and it is possible that the signals seen in the Cb-SNc simply reflect the movements associated with licking and are not related to reward. To disambiguate these relationships, we compared the signals with the licking frequency. Although, as expected, mice increased their licking frequency when they received water and licked for a longer time when they were given a larger volume of water, the differences in the Cb-SNc signals with different rewards did not appear to be fully accounted for by the differences in licking (Figure 8B).

A caveat of these set of experiments was that the mice were continuously licking, although at a lower rate, even prior to receiving reward. This caveat complicated the analysis of a direct correlation between licking and Cb-SNc signals. To remedy this issue, in a different group of animals we used a modified Pavlovian task in which animals were trained to stop licking for at least 3 seconds prior to initiating the trial (Figure 8D, E). Moreover, in addition to expressing GCaMP7 in Cb-SNc axons of these animals, we also expressed jRGECO1 in the SNc dopaminergic neurons by using a Cre-dependent version of the calcium sensor in DAT-Cre mice. By bilaterally placing fiber optics in the SNc, and in the dorsolateral striatum (DLS), we were able to simultaneously monitor changes in the activity of Cb-SNc axons, the cell bodies of dopaminergic neurons within the SNc, and the SNc dopaminergic axons in the DLS as the mice performed the task (Figure 8D, Figure S5). In these set of experiments the mice randomly received either tap water as Regular reward, sweet water as High reward, or no reward at all (Omitted). As in the Pavlovian experiments discussed above, the Cb-SNc axons showed a clear increase in activity when the mice received a reward, with High reward producing a larger signal compared with Regular reward, despite similar licking rates. A much smaller signal, as well as a lower licking rate, was also seen when reward was omitted. These data clearly indicated that the amplitude of the Cb-SNc signals is not simply a scaled function of the licking rate, and that these signals reflect the hedonic value of the reward. Specifically, the licking rate in response to Regular and High rewards were statistically indistinguishable for a 3 s interval following the reward, whereas the Cb-SNc signals were significantly higher for High reward (p<0.001 for each 1 s time bin using post-hoc Holm-Sidak test; following one-way RM-ANOVA, p<0.001, F=12.35).

In these experiments there was a clear increase in the signals recorded from the cell bodies of the SNc dopaminergic neurons and their axonal projections in the DLS, as the animals performed the task (Figure 8E). The amplitude of these signals was also higher in High reward compared to Regular reward trials, with the omission trials producing a transient increase in the signal followed by an undershoot below the baseline. It is noteworthy that the signals arising from the SNc neurons, and those recorded from the axonal projections of the SNc to the DLS were not identical. This could be because the calcium signal in the soma is correlated with both action potentials and subthreshold depolarization, whereas the signal in the DLS corresponds primarily to the action potentials that have successfully propagated along the axon from the SNc to the DLS. Alternatively, axons in the DLS constitute only a subset of projections from the SNc, and the signals sent to different parts of the striatum might not always be the same.

## Discussion

Building on a suggestive body of work accumulated during the past several decades ^11–18^, here for the first time we show that the cerebellum can rapidly modulate the activity of neurons in SNc through an excitatory, monosynaptic pathway. Although both the cerebellum and the basal ganglia are engaged in motor coordination and non-motor behaviors, each structure is thought to have unique features that make it more suitable for specific tasks and computations. The monosynaptic pathway described here might serve as a medium through which the cerebellum updates the basal ganglia with cerebellar-specific, possibly time sensitive, information. In agreement with this hypothesis, we found that the Cb-SNc pathway was bilaterally activated just prior to movement initiation. We also found that activation of the Cb-SNc rapidly increased both the striatal dopamine levels, and the probability that the mice ambulated. Given the modulatory influence of SNc dopaminergic neurons on striatal activity, our findings advance intriguing possibilities regarding cerebellar modulation of the basal ganglia to control motor function, and non-motor behaviors. We also found the Cb-SNc contained information regarding the value of reward, which SNc might deploy to adjust movement vigor, or might use for reward processing per se.

The cerebellum receives a staggeringly wide range of sensory and cortical information and each of its cardinal neurons, the Purkinje cells, integrate over 150,000 synaptic inputs containing this information ^33, 34^. The architecture and neuronal circuitry of the cerebellum serve as an ideal computational platform to integrate this vast information and predict the contraction of different muscles in order to accurately position the various body parts during a complex motor task ^35^. For this reason, it is thought that the cerebellum maintains both forward and inverse internal models of the dynamics of the musculoskeletal system ^36–44^, which enable it to continuously update its predictions to allow for the faithful accomplishment of an intended movement. This unique ability to rapidly make computational predictions distinguishes the cerebellum from the basal ganglia, and might form the basis for the cerebellar-specific time sensitive information that the cerebellum conveys to the basal ganglia.

The time sensitive cerebellar information can be conveyed to the basal ganglia by two pathways. A relatively fast, disynaptic pathway routed through the intralaminar thalamic nuclei allow the cerebellum to modulate the activity of the striatum, the input nuclei of the basal ganglia ^6, 7, 10^. The Cb-SNc pathway described here constitutes the second pathway. Given what is known about the role of SNc in motor control, we propose that the cerebellum might provide these nuclei with information related to movement initiation. The function of the SNc is perhaps best appreciated when one considers its dysfunction in Parkinson’s disease, a motor disorder which is caused by the loss of the dopaminergic neurons in these nuclei. A cardinal feature of Parkinson’s disease is the difficulty of the patients in initiating movements; patients with Parkinson’s disease have a characteristic long delay from the time they resolve to move, to when they actually initiate movement ^45–48^. It is commonly accepted that the reduction in dopamine in the basal ganglia is the major cause of the motor symptoms associated with Parkinson’s disease, as therapeutic approaches that increase dopamine levels are very effective in alleviating the symptoms early in the disease ^49–52^. Thus, dopamine release in the basal ganglia is required for prompt initiation of movements. Recent studies have shown a rapid increase in the activity of dopamine neurons in SNc immediately preceding movement onset ^27–29^, suggesting a role for dopamine in internally-driven movement initiation. Curiously, it has been known for a long time, dating back to observations made by Gordon Holmes, that patients with cerebellar damage have a delay in initiating movements ^53–56^ not unlike the delay in movement initiation seen in patients suffering from Parkinson’s disease. Moreover, inhibiting the activity of the cerebellar nuclei slows reaction times in motor tasks ^57–60^, while activating the cerebellar nuclei by inhibition of Purkinje cells can induce movement ^61^, suggesting that the cerebellum is involved in the timing of initiation of movements. Given that in many cases the activity of neurons in the cerebellar nuclei precedes movement onset ^62–66^, it is plausible that the cerebellum predicts the timing of movement initiation by integrating its cortical and sensory inputs, and conveys these predictions to the SNc via the direct pathway shown here. By doing so, cerebellum can help facilitate the release of a burst of dopamine in the striatum immediately preceding movement onset. In other words, the cerebellum may be the structure that predicts when a movement needs to be made and conveys this information through the monosynaptic pathway to the SNc, which can then release dopamine in the striatum to facilitate the movement. This hypothesis is supported by our observations that activation of the Cb-SNc axons rapidly increases the dopamine levels in the striatum, and that when the animal ambulates or pushes a lever, the pathway is activated just prior to movement initiation.

A wealth of information suggests that SNc and striatal dopamine levels contribute to movement vigor, in addition to movement initiation ^27–32^. Given that the activity of Cb-SNc pathway contains information related to reward value, it is plausible that the cerebellum conveys this information to the SNc to contribute to determination of the movement vigor by promoting greater or less dopamine release for different reward expectations. One caveat of this hypothesis is that our data did not show an increase in the rate of licking that was unambiguously proportional to, and time locked with, the larger Cb-SNc signals when the mice consumed the higher value sweet water reward. One explanation for this lack of scaling, however, might be that licking is a stereotyped motor behavior with a limited dynamic range and the frequency of movements might have already reached its maximum.

If the cerebellum contributes to movement initiation and vigor under normal conditions by promoting release of dopamine by SNc, it is possible that the brain may take advantage of this pathway to compensate for dopamine dysfunction in movement disorders such as Parkinson’s disease. As mentioned above, Parkinson’s disease is caused by loss of the dopaminergic neurons in the SNc, leading to depletion of dopamine in the striatum, and thereby precipitating a number of motor symptoms, including a difficulty in rapidly initiating a movement ^67^. It is plausible that under pathological conditions when there are fewer dopaminergic neurons in the SNc, the cerebellum might increase its input to this structure in order to normalize dopamine levels. In support of this hypothesis, an increase in cerebellar activity, as measured by the expression of immediate early genes, has been observed in dopamine depletion models of Parkinson’s disease in mice ^68^ and monkeys ^69, 70^. Consistent with these data, to complete simple motor tasks, patients with Parkinson’s disease have been shown to recruit the cerebellum more than heathy control subjects ^71–74^, thereby providing support for the notion that the altered activation pattern of the cerebellum in Parkinson’s disease is a mechanism of compensating for dopaminergic dysfunction ^71, 72^. We cannot rule out the alternative possibility, however, that irregular cerebellar activity may be a contributing factor to some of the motor symptoms associated with Parkinson’s disease ^75^.

If the direct Cb-SNc pathway is used by the brain as a compensatory strategy in Parkinson’s disease to normalize dopamine levels in the basal ganglia, a number of testable predictions can be made. First, if the cerebellum attempts to compensate for dopamine loss in Parkinson’s disease, then increased cerebellar activity should be present prior to the complete degeneration of dopamine neurons in SNc. This prediction is supported by the finding that hyperactivity of the cerebellum is a feature of the early stages of Parkinson’s disease ^76^. The second prediction is that increasing the activity of this pathway might be therapeutic, provided that the cerebellum is not already optimally engaged as a compensatory mechanism. Neuromodulation, through transcranial magnetic stimulation (TMS), transcranial direct current stimulation (tDCS), or deep brain stimulation (DBS) of the cerebellar nuclei might be of value in patients with Parkinson’s disease. Both cerebellar targeted TMS and tDCS, but not DBS, have been tested as a means of reducing levodopa-induced dyskinesia and tremor in Parkinson’s disease, but their effects on other symptoms, namely akinesia, have not been examined ^77–80^, although one pilot study showed a slight, but significant, reduction in the time it took patients to perform a voluntary motor task ^81^. While cerebellar DBS has been used in the treatment of ataxia ^82^ and dystonia^83^, it has not been tested in patients with Parkinson’s disease.

Given the prominent roles of both the cerebellum and SNc in motor coordination, the Cb-SNc pathway undoubtedly contributes in some way to motor control, but it is also possible that it may have a complementary function in non-motor behaviors. A wealth of evidence supports the hypothesis that phasic dopamine signaling in the striatum encodes reward prediction error, making it essential for reinforcement learning and reward-seeking behavior ^84–86^. In recent years a growing body of evidence has amassed that also supports a role for the cerebellum in the reward and addiction pathway ^87–95^. Intriguingly, several recent studies have shown that the cerebellum contains reward related information ^88, 96–103^. In addition to the recently described direct projection of cerebellum to one of the major reward processing brain regions, the ventral tegmental area ^104^, the Cb-SNc pathway described here may also serve as a medium through which information about reward and other non-motor behaviors might be conveyed from the cerebellum to the basal ganglia. In support of this hypothesis, in our simple Pavlovian tasks, we found that the activity of the Cb-SNc axons contained information that correlated with the presence and value of the reward.

The cerebellum and basal ganglia are subcortical structures with well-established functions in the motor system and increasingly important roles in reward-based learning and other non-motor behaviors. Understanding how rapid modulation of the SNc by the cerebellar nuclei enables the cerebellum to alter dopamine input to the striatum and ultimately affect basal ganglia output would no doubt shed light on the circuits and mechanisms responsible for generating voluntary movements, and reward-based learning under physiological conditions. Further scrutiny of this pathway is also likely to unearth its role in generating motor symptoms in various disease states and may provide new avenues of therapeutic exploration.

## Acknowledgments

The work was funded, in part, by R01MH115604 and R01DA044761. We thank the members of the Khodakhah lab, and Dr. Saleem Nicola for helpful discussions and feedback. dLight used for dopamine measurements was a kind gift of Dr. Lin Tian.

## Author contributions

S.W. and R.K.B. performed the *in vivo* electrophysiology experiments and post-hoc histology. S.W. performed the *in vivo* electrophysiological recordings and post-hoc histology. M.O. performed the tracing experiments. J.V. performed the dopamine measurements experiments and the fiber photometry experiments in head-restrained animals on the wheel, and post-hoc histology was performed by M.O. J.Y. performed fiber photometry experiments with the lever-manipulation and Pavlovian tasks, and post-hoc histology was performed by L.K. F.N. supervised and contributed to the analysis of the behavioral experiments. K.K. supervised all aspects of the research, and secured funding.

## STAR Methods

### RESOURCE AVAILABILITY

#### Lead contact

Further information and requests for resources and reagents should be directed to and will be fulfilled by the lead contact, Kamran Khodakhah.

### MATERIALS AVAILABILITY

This study did not generate new unique reagents.

This study did not generate new mouse lines.

### Experimental model and subject details

#### Mice

For this study, both female and male mice 6-20 weeks of age were used, except for *in vitro* experiments where the animals were juveniles, P21. Wild-type C57Bl/6, RCE:loxP and DAT^IRES*Cre*^ mice were obtained from Jackson Laboratory (JAX#000664, JAX#032037 and JAX#006660 respectively). All experiments have been approved by the Institutional Animal Care and Use Committee (IACUC) of Albert Einstein College of Medicine. Mice were housed in a 12:12 reverse light:dark cycle with food and water available ad libitum, except for the cohort used for lever-manipulation and Pavlovian tasks who were water restricted (described below).

#### Stereotaxic surgery for AAV injection and fiber optic implantation

All stereotaxic surgeries were performed with mice under isoflurane anesthesia (4-5% for initial sedation, and then maintained at 1.5-2%) with regular monitoring for stable respiratory rate and absence of tail or toe pinch response. Mice were placed in the stereotactic apparatus and a heating pad was used to prevent hypothermia. The skull hair was shaved, a midline incision was made, and the surface of the skull was cleaned of all connective tissue. Craniotomies were made using a drill over the site of injection and/or location of fiber optic placement using coordinate from the Mouse brain Atlas (Franklin and Paxinos, 2008). DV coordinate is referenced to the surface of the brain for all injections (except for Operant/ Pavlovian tasks experiments where DV is referenced to Bregma). Injections were performed using a Microinjector (QSI, Stoelting) using a 10 µl Hamilton syringe (30 Ga), or a Nanoinjector (Nanoject III, Drumond), using glass pipettes. Injections were made by creating a 50 µm pocket below the DV coordinate. After the injection, the syringe or the glass pipette was left in place for at least 5 minutes before retracting it. For Fiber Photometry recordings, Fiber optics (200 µm diameter, 0.37 NA, Neurophotometrics, CA, USA) were implanted using a stereotaxic cannula holder (XCL, ThorLabs), and Charisma® (Heraeus Kulzer) was used to secure the fibers to the skull. For head-restrained electrophysiology/fiber photometry recordings, a metal bracket was implanted on the top of the skull using Charisma. In all cases, a final layer of dental cement was added to the top of the skull to secure fibers and brackets. All animals were treated with the analgesic Flunixin (2.5 mg/kg) every 12 hours for 48 hours following surgery. Mice were allowed to recover for at least 3 weeks after surgery, to allow ample time for both recoveries, and for viral expression before experiments.

Details of viruses, volume and coordinates used in each experiment are described below: For *in vivo* electrophysiology a volume of 0.6 µl of AAV1.hSyn.ChR2 (H134R)-eYFP.WPRE.hGH (ChR2, Addgene 26973P, Penn Vector Core) was injected into the cerebellar nuclei (coordinates: -6.00 AP, ±2.3 ML, -2.3 DV and -6.00 AP, ±1.25 ML, and -2.2 DV) at a rate of 0.1 µl/min. A craniotomy approximately 1 mm in diameter was drilled over the SNc on each side of the skull (-2.92 to – 3.64 mm AP; ±0.5 mm – 1.5 mm ML) for neuronal recordings. The recording sites were covered with a silicone adhesive (KWIK-SIL, WPI).

For *in vitro* electrophysiology a volume of 0.4-0.6 µl of AAV1.hSyn.ChR2 (H134R)-eYFP.WPRE.hGH (Addgene 26973P, Penn Vector Core) or AAV9.hSyn.hChR2(H134R)-eYFP.WPRE.hGH (Addgene 26973P, Penn Vector Core) was injected into the cerebellar nuclei (coordinates: -6.00 AP, ±2.3 ML, -2.2 DV and -6.00 AP, ±1.25 ML, and -2.1 DV) at a rate of 0.1 µl/min.

Anatomical tracing experiments: For AAV-mediated anterograde transsynaptic tracing, 0.6 µl of AAV1.hSyn.CreWPRE.hGH (Addgene #105553) was injected into the cerebellar nuclei (coordinates: -6.0 AP, ±2.3 ML, -2.3 DV and -6.0 AP, ±1.25 ML and -2.2 DV) at a rate of 0.1 µl/min. For retrograde tracing, 4 x 50 nl of a mix of AAVrg.pmSyn1.EBFP-Cre (Addgene #51507) and AAV8.Syn.Chronos.tdTomato (Addgene #62726) in a ratio of 4:1 was injected in the SNc (coordinates: -3.16 AP, ±1.4 ML and -3.85 DV) at a rate of 0.1 µl/min. AAV8-tdTomato was used to label the site of injection. Animals were perfused 4 weeks after surgery for histological examination.

For dopamine measurements, AAV1.hSyn.ChR2 (H134R)-eYFP.WPRE.hGH (Addgene #26973P) was injected bilaterally at coordinates: -6.0 AP, ± 2.3 ML, -2.0 DV for Dentate nucleus, -6.24 AP, ± 1.5 ML, -2.0 DV for Interposed nucleus, and -6.37 AP, ± 0.8 ML, -2.0 DV for Fastigial nucleus (0.4 µl per injection site). The dopamine sensor dLight1.1 was injected in the DLS (AAV9.syn.dLight1.1 at coordinates: +0.25 AP, +2.5 ML, -2.25 DV, volume of 0.4 µl). A fiber optic was implanted unilaterally to target the right SNc (coordinates -3.0 AP, +1.5 ML, - 3.6 DV), and the right DLS (at a DV 100 µm above the site of virus injection). Experiments were performed starting 4 weeks after surgery.

For simultaneous recordings of Cb-SNc axons and SNc neurons in mice on the wheel, the calcium sensor GCaMP7f (AAV9-syn-jGCaMP7f-WPRE, Addgene #104488) was injected bilaterally in the cerebellar nuclei, at coordinates: -6.0 AP, ± 2.3 ML, -2.0 DV for Dentate, -6.24 AP, ± 1.5 ML, -2.0 DV for Interposed, and -6.37 AP, ± 0.8 ML, -2.0 DV for Fastigial (0.4 µl per injection site). In addition, the red-shifted calcium sensor jRGECO1a (AAV1-Syn-NES-jRGECO1a-WPRE-SV40, Addgene #100854) was injected in the SNc (coordinates: -3.0 AP, ±1.5 ML, -3.6 DV, 400 nL per injection), and Fiber optics were implanted bilaterally in the SNc 100 µm above the site of virus injection. Experiments were performed starting 4 weeks after surgery.

For fiber photometry experiments measuring calcium activity in SNc in the Operant/Pavlovian tasks, mice were injected with AAV9-syn-jGCaMP7f-WPRE (Addgene, 104488) in the DCN (coordinates: -6.00 AP, ±2.30 ML, -3.50 DV and -6.30 AP, ±1.25 AP, -3.20 DV, 0.6 µl per injection site) and the Fiber optic was implanted in the SNc (coordinates: -3.00 AP, ±1.50 ML, -4.20 DV).

For simultaneous fiber photometry recordings from Cb-SNc axons, SNc.DA-neurons, and SNc.DA. DLS axons in the Pavlovian tasks, mice were injected with 0.6 µl of AAV9-syn-jGCaMP7f-WPRE (Addgene, 104488) in the DCN (coordinates: -6.00 AP, ±2.30 ML, -3.50 DV and -6.30 AP, ±1.25 AP, -3.20 DV), and with AAV1-Syn-Flex.NES-jRGECO1a-WPRE-SV40 (Addgene, 100853) targeting the SNc (-3.00 AP, ±1.50 ML, -4.20 DV). Fiber optics were implanted in the SNc (-3.00 AP, ±1.50 ML, -4.20 DV) and the DLS (+0.25 AP, ±2.50 ML. -3.20 DV).

#### *In vivo* electrophysiology

*In vivo* extracellular single unit recordings were made using an optrode to record the neurons in SNc while optogenetically stimulating ChR2-expressing cerebellar fibers close by. Optrodes were constructed by attaching a 105 µm fiber optic (Multimode Fiber, 0.22 NA, High-OH 105 µm Core, ThorLabs, FG105UCA) to a tungsten electrode (WPI, TM33B20, shaft diameter = 0.256 mm) with superglue and epoxy. The tip of the electrode was positioned less than 250 µm below the end of the fiber optic. The optrode was then coated with Vybrant^TM^ DiI dye (Invitrogen, V22889) for post-hoc confirmation of recording sites. To coat the optrode, it was dipped 10 times into an Eppendorf tube filled with Vybrant dye and then allowed to dry for 20-30 minutes. This process was repeated 3 times. At the time of recording, the craniotomy over the SNc was uncovered and the surface of the brain was detected by touching it with the tip of the electrode. All depths are measured with respect to the surface of the brain. The well around the craniotomy was filled with sterile saline, in which a reference wire was placed. The optrode was then lowered into the brain to the depth of SNc, according to the Paxinos Mouse brain Atlas ^105^.

Once a single unit was isolated, 1 ms pulses of light were applied at 5 second intervals with a 447 nm laser (OEM laser Systems, Inc). Varying light intensities were tested to achieve a minimum and maximum response. Cells that did not respond to the maximum intensity of light tested, (greater than 10 mW) were considered non-responding cells. In most cases, 1 mW – 2 mW of light was needed to evoke a response from a responsive cell, and this intensity was used to measure the response of the cell to cerebellar stimulation. The response of an SNc neuron to a train of stimuli was measured by activating cerebellar fibers with 5 pulses of light, each 1ms in duration, at a frequency of 20 Hz, a frequency that ChR2 can faithfully follow ^106^.

The signal of an isolated cell was amplified 2000x with a custom-made amplifier and digitized at 20 kHz. Data was acquired using custom-written software in LabVIEW (National Instruments), and single spikes were sorted using Plexon® Offline Sorter™. Analysis was performed with custom-written software in MATLAB (Mathworks). The time interval including the 5 seconds before a stimulus and the 5 seconds after the stimulus was divided into 1 ms bins, and the number of spikes in each bin was counted. A histogram of this data was plotted. The presence of an excitatory response, and its latency were determined by identifying the first bin after the stimulus where the number of spikes were three standard deviations greater that the average number of baseline spikes per bin. The duration of the excitatory response was determined as the time from the onset of the response to the first bin were the number of spikes per bin returned to within three standard deviation of the baseline firing rate. The number of extra spikes evoked by a stimulus was calculated by subtracting the number of spikes expected to be included in the excitatory response period based on the baseline firing rate of the cell from the number of spikes observed in the excitatory response period. For trains of stimuli, the extra spikes evoked by all additional pulses were normalized to the extra spikes evoked by the first pulse. The presence of an Inhibitory responses was determined by identifying the presence of three or more consecutive bins after the stimulus where the number of observed spikes was less than 1 standard deviation of the expected baseline spikes.

#### *In vitro* electrophysiology

Mice were anesthetized with isoflurane, transcardially perfused with NDMG aCSF ^107^ (in mM: NMDG 93, KCl 2.5, NaH_2_PO_4_ 1.2, NaHCO_3_ 30, Glucose 25, HEPES 20, Na-ascorbate 5, Na-pyruvate 3, Thiourea 2, MgSO_4_ 10, CaCl_2_ 0.5), and decapitated. The brain was quickly removed and placed in ice cold NMDG aCSF. The brain was then secured with super glue to a metal platform and placed in a bath of NMDG aCSF at 2-4°C. Horizontal slices 250 µm thick of SNc were made from the ventral side of the brain on a vibrating microtome (Campden Instruments Model 7000smz-2). Slices were incubated in NMDG aCSF at 34°C for less than 15 minutes and then placed in standard aCSF (in mM: 125 NaCl, 2.5 KCl, 26 NaHCO3, 1.25 NaH_2_PO_4_, 1 MgCl_2_, 2 CaCl_2_, and 11 glucose) at room temperature for at least an hour.

Slices were used up to 6 hours after slicing. All solutions used during slicing were continuously bubbled with 95% oxygen:5% carbon dioxide.

Slices were transferred to a recording chamber continuously perfused with oxygenated aCSF at a minimum rate of 1.5-2 mL/min and maintained at a temperature of 32-35°C.

Recording electrodes were made from filamented borosilicate glass (Sutter BF150-110-10HP), pulled to a resistance of 2.5-4 MΩ (Sutter Model P-97), and filled with internal solution containing (in mM) 70 Cs-gluconate, 10 CsF, 20 CsCl, 10 EGTA, 10 HEPES, 3 Na2ATP, and 2 QX-314. 2 mM Neurobiotin tracer (Vector Laboratories, SP-1120) was also added to the internal solution.

Midbrain neurons were visualized with a 40x objective under oblique infrared illumination (Zeiss). SNc was identified by the presence of large, densely packed cell bodies located anterior and lateral to the medial terminal nucleus of the accessory optic tract (MT), which appears as a round dark spot in horizontal slices under the microscope ^108^. Whole cell recordings were made with a Cairn Optopatch amplifier, and cells were clamped at -60 mV unless otherwise stated. The ChR2-expressing cerebellar axons in the field of view were optogenetically activated through the objective by an LED (455 nm, ThorLabs M455L3). The duration of each light pulse was 1 ms unless otherwise stated. The maximum stimulus intensity measured at the output of the 40x objective did not exceed 5 mW.

Whole cell data was acquired at 10 kHz (PCI-MIO-16XE-10 or 6052E; National Instruments) using custom written software in LabVIEW. Data was analyzed in LabVIEW and MATLAB.

The following compounds were used in the pharmacology experiments: Cyanquixaline (CNQX, Tocris, Fisher 1045, 10 µM), 2,3-dihydroxy-6-nitro-7-sulfamoyl-benzo[f]quinoxaline-2,3-dione (NBQX, Cayman Chemicals, 14914, 10 µM), D-(-)-2-Amino-5-phosophonopentanoic acid (D-APV, Tocris, R&D Systems, 0106, 50 µM), Tetrodotoxin (TTX, Cayman Chemicals, 19084-100, 1 µM), and 4-Aminopyridine (4-AP, Sigma-Aldrich, 278587, 200 µM).

#### Immunohistochemistry

For post-hoc histology, mice were anesthetized with isoflurane and transcardially perfused with phosphate buffered saline (1M PBS) followed by 4% paraformaldehyde (PFA). The brains were dissected and fixed overnight in 4% PFA at 4°C. They were then rehydrated with 15% sucrose for 24 hours and then with 30% sucrose for 24-48 hours, and rapidly frozen in Optimal Cutting Temperature Compound (OCT) on dry ice. The brain was cut on a cryostat (Leica CM3050 S) into 30 µm sections for all experiments except for tracing experiments, where 25 µm sections were used. Sections were blocked with 5% goat serum (Gibco) in 0.3% PBS-Triton for 1 hour. Incubation with primary antibodies was performed overnight at 4°C in blocking solution, followed by secondary antibodies incubation for 1 hour in blocking solution. The following antibodies were used: anti tyrosine hydroxylase (TH, rabbit, Millipore, AB152), anti-GFP (chicken, Abcam, ab13970), anti-RFP (rabbit, Rockland, 600-401-379), anti-TH (mouse, Millipore, MAB318), anti-RFP (guinea pig, Synaptic Systems, 390 004), Alexa Fluor 568 (goat anti-rabbit, Molecular Probes A11011), Alexa Fluor Plus 647 (goat anti-rabbit, Invitrogen Life Technologies, A32733), Alexa Fluor 488 (goat anti-chicken, Molecular Probes, A11039), Alexa Fluor Plus 594 (goat anti-rabbit, Invitrogen Life Technologies, A32740), Alexa Fluor Plus 647 (goat anti-mouse, Molecular Probes, A21240), Alexa Fluor 568 (goat anti-guinea pig, Molecular Probes, A11075). Nuclei were labeled with DAPI (Hoescht 33342, Invitrogen Life Technologies, H3570). Neurobiotin was detected with Streptavidin conjugated to Alexa 568 (2 ug/mL, Invitrogen Life Technologies, S11226). Recording sites were determined by the presence of Vibrant® DiI dye (Invitrogen V-22889) fluorescence, or by a lesion in the tissue made by the optrode (for details of antibodies/dyes used in each experiment see Table S1).

Slides were mounted with Fluoromont-G (Southern Biotech) and covered for further imaging. Images of sections were captured using either a standard fluorescent microscope (Zeiss) at 40x magnification, a confocal microscope (Zeiss LSM880 Airyscan with super-resolution) at 63x magnification, or a slide scanner microscope (Zeiss Axio Scan.Z1) at 20X magnification.

#### Anatomy data analysis

Quantification of the number of DCN-recipient cells in the SNc was performed in 25 µm thick serial sections of the entire midbrain every 75 µm. For antero-posterior (A-P) analysis, the total number of GFP cells was calculated in each section in the SNc and matched with a bregma position according to the Paxinos Mouse Brain Atlas ^105^. Since anterograde tracing was performed bilateral in the DCN, data from each hemisphere in the SNc was combined and a mean value was estimated for the analysis in each section, respectively. Counting of GFP cells was performed in Fiji (ImageJ, NIH).

Counting of SNc projecting neurons in the DCN (input cells) was performed in 25 µm thick serial cerebellar sections every 75 µm. The number of GFP input cells in each nucleus of the DCN was calculated in each mouse. Percentage of cells in each DCN nuclei was calculated from the total number of GFP cells in the DCN of each mouse. Total area of each DCN was measured using Fiji, as well as the number of input cells to calculate the density.

Mapping of neurons in the SNc and DCN that comprise the Cb-SNc was performed using the Paxinos Mouse Brain Atlas ^105^ as reference, using the data obtained from anterograde and retrograde tracing experiments. Brain Atlas schematics shown in the figures are approximately 120 microns apart. Each schematic was populated by mapping the labeled neurons identified in the third and sixth consecutive 25-micron thick serial section that best corresponded with the Atlas coordinates.

#### Head-restrain treadmill

This apparatus was described by Heiney et al. (Heiney, Wohl, Chettih, Ruffolo, & Medina, 2014). The treadmill is made of a foam cylinder with bearings holding a metal axle that is secured to a metal frame. The head-bracket is attached to the metal frame using two screws, leaving mice head-restrain on top of the cylinder and free to walk at will. Walking activity was recorded by using a rotary encoder coupled to the cylinder with a rubber band, which analog signal was digitized at 5kHz using the analog-to-digital converter USB6001 (National Instruments, TX, USA) and recorded with the Bonsai software (www.open-ephys.org/bonsai). The rotary encoder signal was calibrated to convert the voltage amplitude to speed (cm/s).

#### Quantification of dLight1.1 dopamine signals

Dopamine fluctuations were recorded using fiber photometry by measuring the fluorescence emitted by a dopamine sensor (AAV9.syn.dLight1.1, a kind gift of Dr. Lin Tian). dLight1.1 was exited with a 470 nm LED via a patch cable (200 µm fiber optic, Docris), and the emitted fluorescence was recorded at 40 FPS. Spontaneous events were identified by eye and aligned to time zero (onset) when the signal went above 3 SD of the baseline (200 ms window). Optogenetic activation of Cb-SNc axons was performed with a 447 nm laser (OEM systems, 1 mW total output), via an optic fiber implanted in SNc, and triggered by a computer running a custom-witen MATLAB script and connected to a digital-to-analog converter USB6001. The stimulation patterns used were 1 ms single pulse (inter pulse interval of 8 s ± 2 s jitter), a short train of 15 1 ms-pulses at 20 Hz (inter train interval of 10 s ± 2 s jitter), and a long train of 50 1 ms-pulses at 20 Hz (inter train interval of 15 s ± 2 s jitter). Each pattern was applied for 3-5 minutes twice per session in random order.

#### Correlation between dopamine signals and time walking

Dopamine signals where quantified as the integral of the fluorescence trace in a time window (400 ms for Single Pulse, 1.4 s for Short Train, and 3.4 s for Long Train) multiplied by the number of stimulations performed per minute (7.5 Single Pulse, 6 Short Train, 4 Long Train) and normalized by the average value.

We quantified the percentage of time walking as the total time mice spent moving (detected as a non-zero signal in wheel’s speed) divided by the total time mice were stimulated with Single Pulses (4 min, inter stimulus interval 8 s), Short Trains (2 mins, inter stimulus interval 10 s), or Long Trains (2 min, inter stimulus interval 15 s). The correlation was performed using 3 data points (Single Stim, Short and Long Train) per mouse (N=3).

#### Head-Restricted Pavlovian and Operant Task

After recovery from the stereotaxic surgery (>3 weeks), mice were water deprived. Mice body weight and general health was monitored daily. Typically, the mice obtained their full daily intake of water after completion of the reward task experiments. However, when necessary, 0.5∼1.0 mL water was provided to the mice to maintain >85% of the original body weight. Food was available in the home cage ad libitum. The head-restricted Pavlovian and operant tasks were performed using the Task Forcer equipment (O’hara, Tokyo, Japan).

#### Operant Lever-Manipulation Task

To monitor SNc-projecting cerebellar activity during operant behaviors, we used a lever-manipulation task. During the task session, the mouse’s head was fixed by holding the metal bracket. A lever was introduced to the right hand. The lever was movable horizontally (full-range 2 mm; positions defined as follows: 1.4 a.u. for front end, 0.0 a.u. for the rear end). The mouse manipulates the lever and gets water from a spout located ahead of the mouth after performing correct lever manipulation.

At the beginning of the task trial, the lever was automatically held at the rear end for 3.0 s. After completing the automatic 3.0 s lever-hold, the mouse was allowed to move the lever to the front side. When the mouse moved the lever to the front end of the movable range within 5.0 s, the lever was held at the front end, and the 0.3 s light cue was delivered with 1.0 ∼ 2.0 s delay (When the mouse did not move the lever withing 5.0 s, the lever was automatically returned to the rear end and the subsequent trial started). 4.0 μL water was delivered as a reward via the spout located in front of its mouth 1.0 s after the light cue delivery. The lever was automatically returned to the rear end 2.0 s after the reward delivery, and the subsequent trial started. A task session was stopped when the mouse completed 300 trials. %). Lever position and task-event analog signal were recorded via NI-USB 6001 (National Instruments, TX, USA) at a 1000 Hz sampling rate.

#### Pavlovian Random Reward Task without Licking Interruption

To monitor SNc-projecting cerebellar activity related to reward evaluation, we utilized a Pavlovian task. The mouse’s head was fixed as same as the above operant task during a session. At first, a house lamp (conditional stimulus: CS) was turned on for 0.1 s every 6.0 ∼ 10.0 s. 1.0 s after the CS, the various liquid reward was randomly delivered to the mouse via a spout. Each reward was 4 µL tap water (Regular; 70%), 4 µL 0.1% saccharine water (High; 10%), three shots of the 4µL tap water (3xRegular; 10%), and no-reward (Omission; 10%). Licking during the session was recorded using an Infrared night vision high-speed camera (Shenzhen Ailipu Technology, Shenzhen, China) at 100 fps. Task-event analog signal was recorded via NI-USB 6001 (National Instruments, TX, USA) at a 1000 Hz sampling rate.

#### Pavlovian Random Reward Task with Licking Interruption

To monitor reward-related SNc-projecting cerebellar activity without the influence of the licking-related activity, we introduced the licking interruption period in a Pavlovian task. Licking during a session was detected as spout vibration and utilized for the task control. The house lamp was turned on for 0.3 s as a conditional stimulus when a head-restrained mouse stayed without any licking for 3.0 ∼ 5.0 s. In this Pavlovian task, the conditional stimulus and various rewards were delivered at the same time. 4 µL tap water (Regular; 33.3%) or 4 µL 0.1% saccharine water (High; 33.3%) or and no-reward (Omission; 33.3%) were randomly delivered. 6.0 ∼ 10.0 s inter-trial interval was introduced when a licking was detected after the conditional stimulus. %). Licking during the session was recorded using an Infrared night vision high-speed camera (Shenzhen Ailipu Technology, Shenzhen, China) at 100 fps. Task-event analog signal was recorded via NI-USB 6001 (National Instruments, TX, USA) at a 1000 Hz sampling rate.

#### Fiber photometry Data Collection and Analysis

Fiber photometry fluorescence signals were collected with a fiber photometry apparatus (FP3001, Neurophotometrics Ltd.) at 40 FPS, with 24 ms integration time per frame. 470 nm and 560 nm LEDs at power intensity of 1.6-3.2 mW/mm^2^ were used to excite green (GCamP7f, dLight1.1) and red-shifted (jRGECO1a) fluorescent sensors, respectively. Recordings were performed either using a single-color stimulation (constant mode) in which a single LED was constantly on, or using a two-colors stimulation (multiplexed mode) in which each LED was active every other frame. Detected fluoresce was recorded using the Bonsai software (www.open-ephys.org/bonsai).

For the dopamine measurements, offline analysis of recorded traces was performed with the IgorPro 9 software (Wavemetrics.com), and consisted in the subtraction of a double exponential, and the generation of stimulus triggered averages.

For the dual-color fiber photometry recordings on the treadmill wheel, offline analysis of recorded traces was performed with IgorPro 9 software (Wavemetrics.com). Events were time-locked to the onset of locomotor activity (defined as a wheel speed 3 time the SD of the baseline signal when the mouse was stationary).

For the operant and Pavlovian tasks, lever-position, task-event analog signal, and video recordings of the mice as they performed the task were collected using Bonsai software(www.open-ephys.org/bonsai). The collected data was analyzed using custom-written routines in MATLAB (Mathworks, CA, USA).

In all cases the signals were corrected for bleaching by subtraction of a fitted double exponential to the raw signals. Z-scored signals were calculated by obtaining the mean and standard deviation of the corrected signals from a baseline period (treadmill baseline was the 4 s period prior to locomotion, Operant task baseline was 1 s to 0.1 s prior to lever movement, and Pavlovian task baseline was 750 to 250 ms prior to cue).

#### Statistics

Statistical analyses were performed using either GraphPad Prism 7 (GraphPad Software), or Mathlab. Unless otherwise stated, data were assessed for normality using Shapiro-Wilk normality test. Normally distributed datasets were statistically compared based on the two-tailed, paired Student’s t-test or one-way ANOVA with Tukey’s correction or one-way repeated ANOVA with Holm-Sidak correction for multiple comparisons.

Unless otherwise stated, for statistical tests alpha was set at 0.05. Variability within datasets is represented by box plots showing the median, interquartile range, and 10–90 percentiles (error bars) or a histogram. Data are reported in text as mean ± S.D. unless otherwise stated.

---

**Figure S1.**
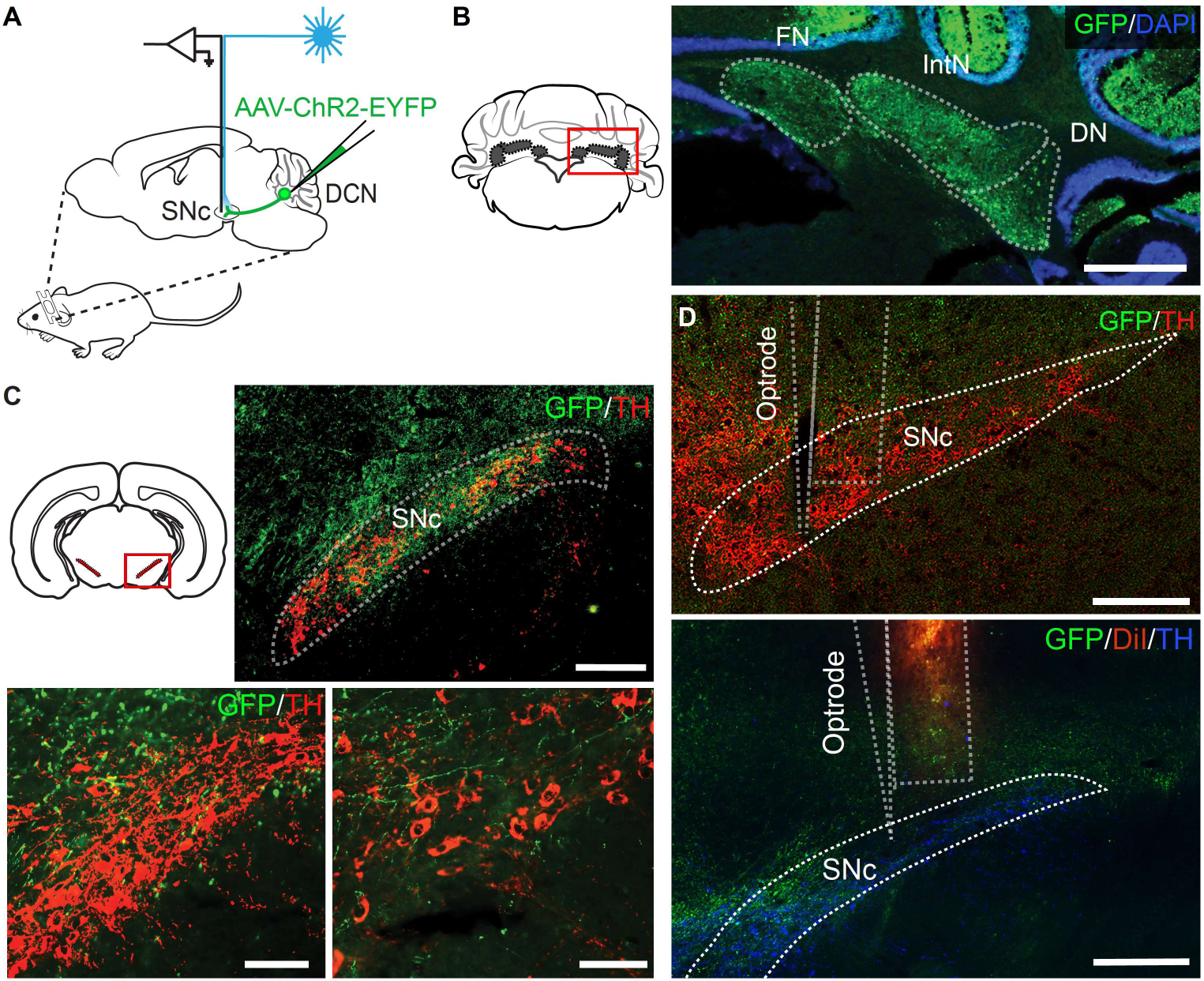
Post-hoc histology of *in vivo* electrophysiology experiments, related to Figure 1. (A) AAV-ChR2-EYPF was injected into the DCN and an optrode was placed in the SNc for *in vivo* recordings. (B) Schematic of DCN region zoom in. Expression of ChR2 was analyzed with GFP (green) in the cerebellar nuclei. Nuclei were visualized with DAPI. All three DCN (FN, IntN and DN) were targeted. Scale bar: 1 mm (C) (Top, left) Schematic of SNc region zoom in. (Top, right) Expression of ChR2 in cerebellar fibers in SNc was visualized with GFP (green). TH staining was performed to delimit the structure and to label for dopaminergic neurons (red). Scale bar: 250 µm. (Bottom) Different examples showing cerebellar axons in SNc. Scale bars: 100 µm. (D) (Top) Example of the lesion in the tissue caused by the optrode in the SNc. (Bottom) To track the position of the optrode, the electrode was labeled with DiI (red). TH is shown in blue. Scale bars: 250 µm.

**Figure S2.**
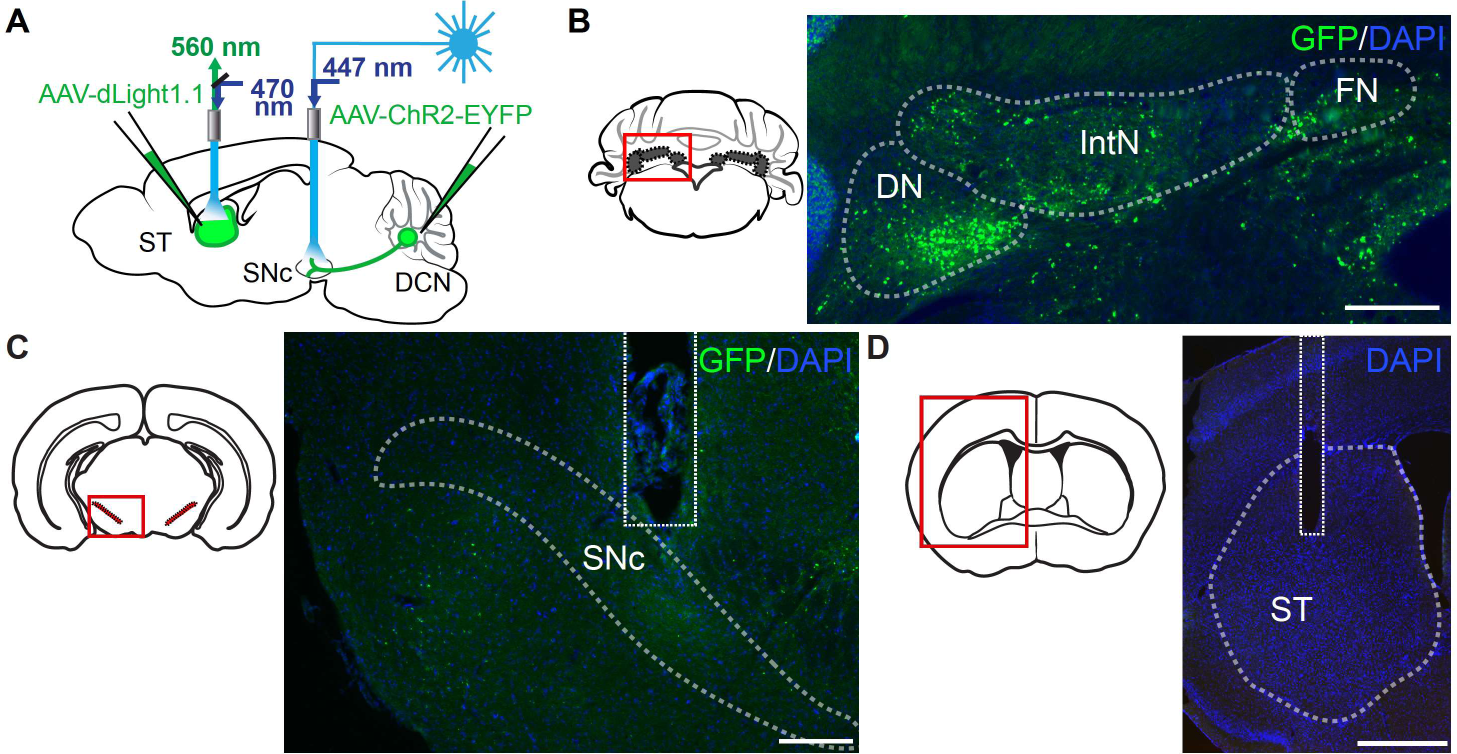
Post-hoc histology of dopamine measurements experiments, related to Figure 6. (A) AAV-ChR2-EYFP was injected in the DCN and AAV-dlight1.1 in the striatum. Fiber optics were implanted in the SNc to stimulate Cb-SNc axons and in the striatum to measure dopamine fluctuations. (B) Schematic of DCN region zoom in. Expression of ChR2 with GFP (green) in the deep cerebellar nuclei. All three nuclei (FN, IntN, DN) were infected. Nuclei were visualized with DAPI in blue. Scale bar: 250 µm. (C) Schematic of SNc region zoom in. Fiber optic location in SNc. Scale bar: 250 µm. (D) Schematic of striatum region zoom in. Fiber optic location in striatum. Scale bar: 1 mm.

**Figure S3.**
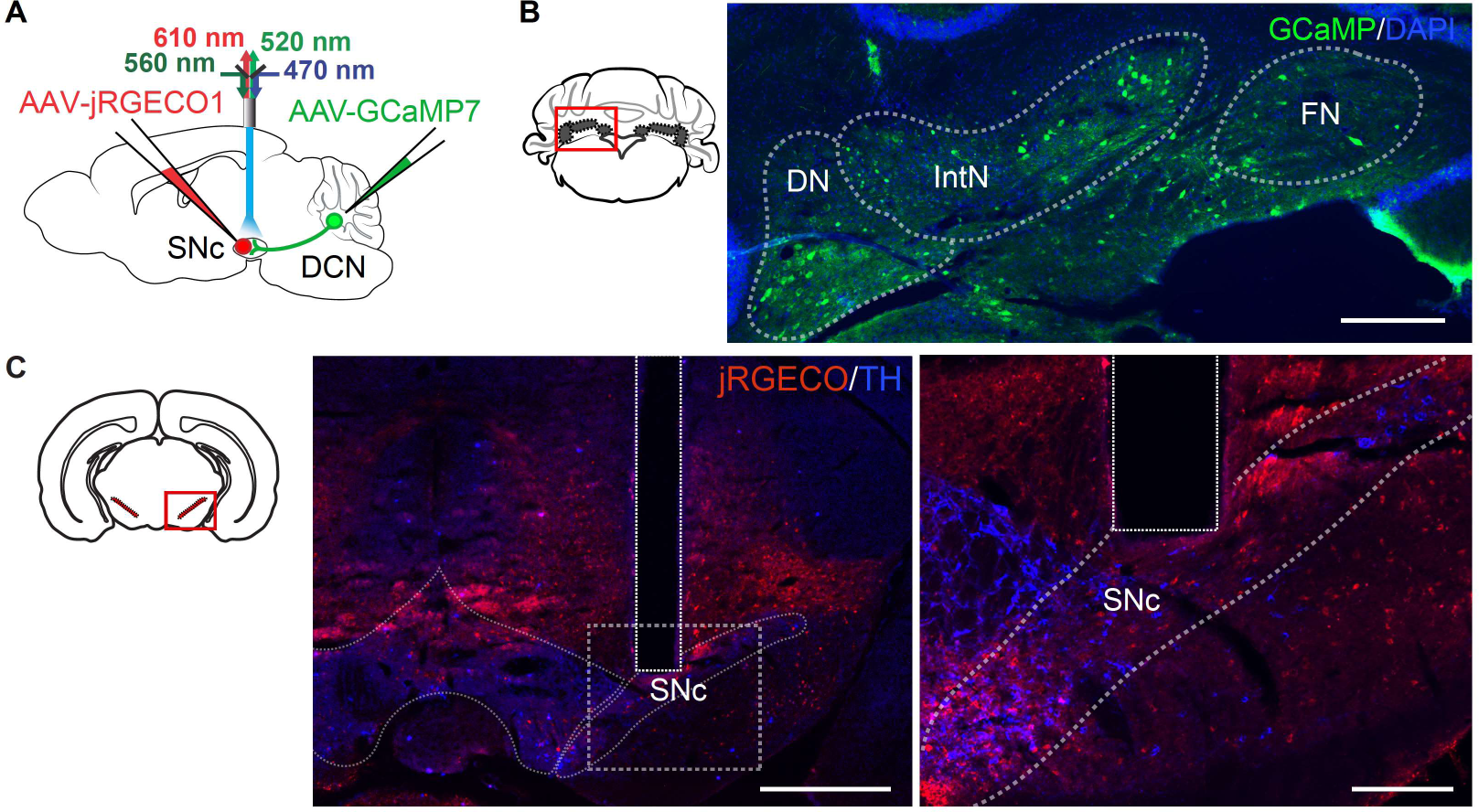
Post-hoc histology of simultaneous fiber photometry recordings in mice on the wheel, related to Figure 7 A-C. (A) AAV-GCaMP7 was injected in the DCN and AAV-jRGECO1 was injected in the SNc. Fiber optic was implanted in the SNc for dual recordings. (B) Schematic of DCN region zoom in. Expression of GCaMP (green) in the deep cerebellar nuclei. All three nuclei were infected (FN, IntN, DN). Nuclei were visualized with DAPI in blue. Scale bar: 250 µm. (C) Schematic of SNc region zoom in. (Left) Expression of jRGECO and fiber location in SNc. Scale bar: 1 mm. (Right) Zoom in of left image showing jRGECO neurons in red. TH was used to delimit SNc area (blue) Fiber optic is located above the SNc. Scale bar: 250 µm.

**Figure S4.**
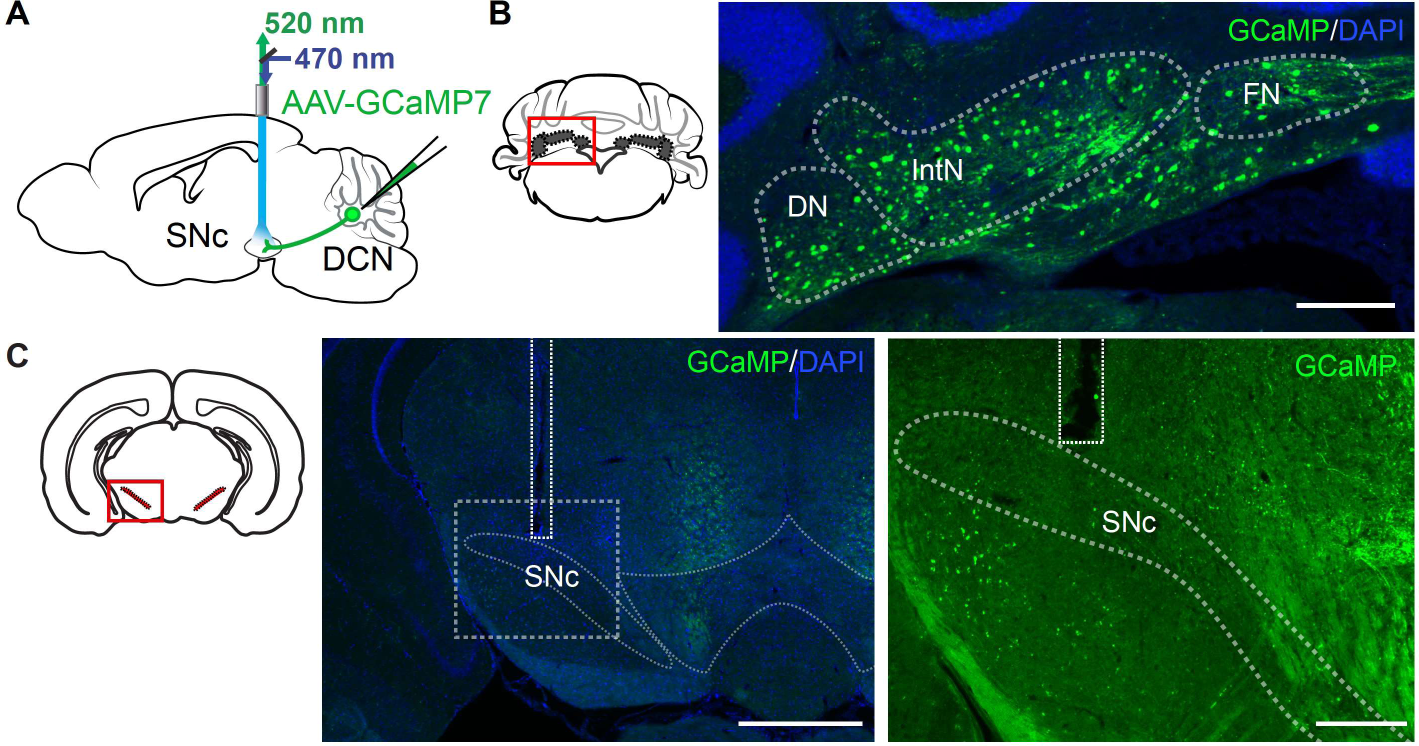
Post-hoc histology of single fiber photometry recordings in mice performing the lever-manipulation and Pavlovian tasks, related to Figures 7 D-E and Figure 8 A-C. (A) AAV-GCaMP7 was injected in the DCN. Fiber optics were implanted in the SNc to measure Cb-SNc axons activity. (B) Schematic of DCN region zoom in. Expression of GCaMP (green) in the deep cerebellar nuclei. All three nuclei (FN, IntN, DN) were infected. Nuclei were visualized with DAPI in blue. Scale bar: 250 µm. (C) Schematic of SNc region zoom in. Fiber optic location in SNc. Scale bar: 250 µm.

**Figure S5.**
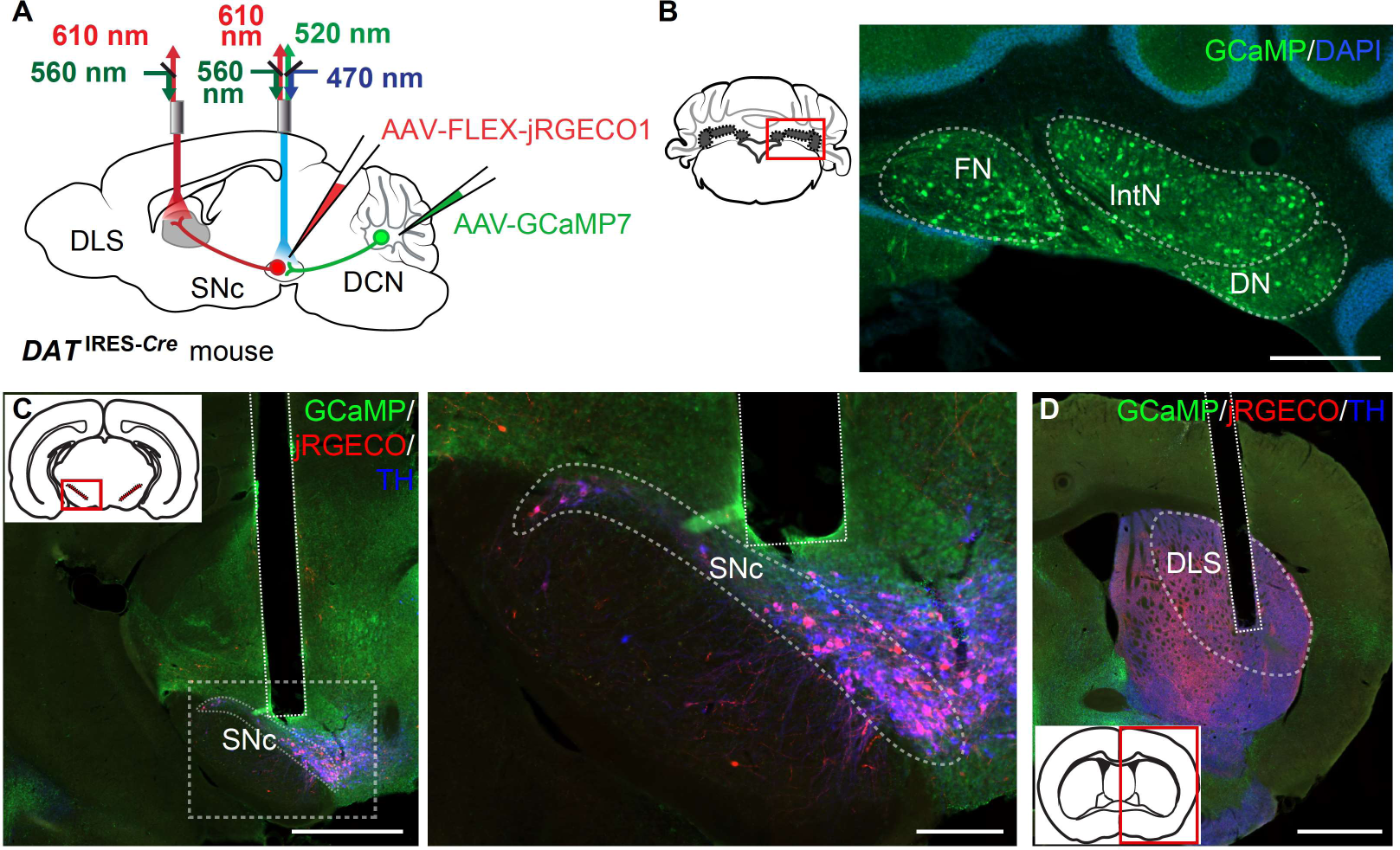
Post-hoc histology of dual fiber photometry recordings in mice performing Pavlovian task, related to Figure 8 D-E. (A) AAV-GCaMP7 was injected in the DCN and AAV-FLEX-jRGECO1 in the SNc of DAT-Cre mice. Fiber optic were implanted in the SNc to measure Cb-SNc axons and SNc-DA neurons activity while fiber in the dorsolateral striatum (DLS) recorded SNc-DA neurons axons. (B) Schematic of DCN region zoom in. Expression of GCaMP7 (green) in the cerebellar nuclei. All three nuclei were infected (FN, IntN, DN). Nuclei were visualized with DAPI in blue. Scale bar: 1 mm. (C) Schematic of SNc region zoom in. (Left) Expression of jRGECO and fiber location in SNc. Scale bar: 1 mm. (Right) Zoom in of left image showing jRGECO neurons in red colocalized with TH (white) and Cb-SNc fibers expressing GCaMP (green). Fiber optic is located above the SNc. Nuclei were visualized with DAPI in blue. Scale bar: 250 µm. (D) Zoom in schematic of striatum. jRGECO expression in SNc axons (red). TH expression is shown to help identify the boundary of the striatum (white). Fiber optic location in DLS. Scale bar: 1 mm

